# A *Trypanosoma brucei* orphan kinesin employs a convergent microtubule organization strategy to complete cytokinesis

**DOI:** 10.1101/2021.11.04.467292

**Authors:** Thomas E. Sladewski, Paul C. Campbell, Neil Billington, Alexandra D’Ordine, Christopher L. de Graffenried

## Abstract

Many single-celled eukaryotes have complex cell morphologies defined by cytoskeletal elements comprising microtubules arranged into higher-order structures. *Trypanosoma brucei* (*T. brucei*) cell polarity is mediated by a parallel array of microtubules that underlie the plasma membrane and define the auger-like shape of the parasite. The subpellicular array must be partitioned and segregated using a microtubule-based mechanism during cell division. We previously identified an orphan kinesin, KLIF, that localizes to the division plane and is essential for the completion of cytokinesis. To gain mechanistic insight into how this novel kinesin functions to complete cleavage furrow ingression, we characterized the biophysical properties of the KLIF motor domain *in vitro*. We found that KLIF is a non-processive dimeric kinesin that dynamically crosslinks microtubules. Microtubules crosslinked in an antiparallel orientation are translocated relative to one another by KLIF, while microtubules crosslinked parallel to one another remain static, resulting in the formation of organized parallel bundles. In addition, we found that KLIF stabilizes the alignment of microtubule plus ends. These features provide a mechanistic understanding for how KLIF functions to form a new pole of aligned microtubule plus ends that defines the shape of the new posterior, which is a unique requirement for the completion of cytokinesis in *T. brucei*.

## Introduction

Many parasites display high degrees of cell polarity and organization that play essential roles in host colonization and subsequent transmission while mitigating the host immune response. In eukaryotic parasites, cell polarity is frequently established by microtubule-containing structures, such as the cortical microtubules in *Toxoplasma gondii* and the ventral disc in *Giardia lamblia* (1, 2). The assembly and maintenance of these structures require unique adaptations to fundamental cellular processes, such as cell division, that must duplicate and position them within highly polarized cell bodies (3, 4). Understanding how cell division pathways have been tuned to account for the specialized morphologies of these organisms, which have evolved along different evolutionary tracks than most well-studied eukaryotes, could provide insight into the range of potential mechanisms available for building a new cell.

Cytokinesis occurs during the last stages of cell division and is responsible for segregating the duplicated organelles to form two new daughter cells. In most highly studied eukaryotes, cytokinesis is driven by the formation of a tension-generating actomyosin ring that provides the force for cleavage furrow ingression at a symmetric point within the cell body (5-7). However, the ubiquity of this mechanism has been called into question by the sequencing of a broader range of eukaryotes, especially among the unicellular organisms that make up the bulk of eukaryotic diversity (8). For example, the protist parasite *Trypanosoma brucei* (*T. brucei*) lacks myosin II (5-7) and does not appear to require actin for cell division in the procyclic form, which suggest that an actomyosin ring is not employed for cytokinesis (5, 9). These differences argue that cytokinesis may be a more mechanistically diverse process than previously considered, which opens the possibility that actomyosin represents a niche approach to an essential cell-cycle event when put in the context of a broader range of eukaryotes.

*T. brucei* is the causative agent of Human African Trypanosomiasis and the related illness nagana in ungulates. The parasite has an extended, corkscrew-like morphology essential for its motility and infectivity that is mediated by a subpellicular array of highly crosslinked microtubules (MTs) underlying the plasma membrane. The MTs comprising the array are arranged with their plus ends concentrated at the cell posterior and minus ends at the cell anterior. The parasite retains its shape throughout cell division, which requires the enlargement and reorganization of the subpellicular array so that the daughter cells can be built from the existing polarized structure (10). Cytokinesis in *T. brucei* occurs via a furrow that ingresses unidirectionally from the anterior end of the cell towards the posterior. Ultrastructural studies have shown that the cleavage furrow initially moves along an in-folded section of the subpellicular array that likely facilitates the rearrangement of the MTs into two discrete MT arrays (11). During the last stages of cytokinesis, a new cell posterior is constructed via an unknown process that brings together a bundle of aligned MT plus ends that define the shape of the new posterior (11, 12).

We previously used proteomic approaches to identify a number of novel proteins that play essential functions during *T. brucei* cytokinesis (13). Among these proteins is a previously unstudied kinesin we named KLIF (Kinesin Localized to the Ingressing Furrow). KLIF localizes to the site of cleavage furrow initiation, which is present at the anterior end of the duplicating cell, and subsequently along the furrow as it ingresses towards the posterior (13). Initial *in vitro* characterization of the KLIF motor domain determined that it moves slowly along MTs and is plus end directed. Depletion of KLIF using RNAi results in cell cycle arrest and an accumulation of cells with partially ingressed cleavage furrows, indicating that the motor is needed to complete the final stages of cytokinesis as the new cell posterior is created.

Kinesins are the only universally conserved class of molecular motors found in eukaryotes (14). While the motor domains show some degree of conservation across kinesin families, their overall domain architectures are highly divergent, suggesting that there are a wide variety of kinesin configurations that transform the force generated by the motor into cellular function. This variety includes the position of the motor domain within the polypeptide chain, the number and location of the other specialized domains involved in protein binding, and the oligomerization state and orientation of the motor heads. A bioinformatic survey of kinesins from a broad range of eukaryotes identified 105 different domain architectures, of which only 28 are highly conserved, suggesting a high degree of specialization at the organismal level. While conserved domain architectures can be used to predict related functions across organisms, KLIF falls into a category of “orphan” or ungrouped kinesins that lack homology to established motor families (14, 15). This makes it difficult to propose a specific function for KLIF in *T. brucei* cytokinesis, where it could be functioning in processes such as the delivery of essential cargo to the furrow using the array MTs as a track or the reorganization of the array MTs to duplicate the subpellicular array.

In this work, we have used *in vitro* approaches to visualize how KLIF interacts with a simplified set of MTs that mimic the *T. brucei* subpellicular array. This approach has revealed that while the sequence of KLIF diverges from any known class of kinesin, its molecular function shares similarities with kinesins that organize MT networks. These data are the first biophysical description of any kinesin in *T. brucei* and provide essential insight into how KLIF functions to complete the final stages of cytokinesis. We show that a minimal KLIF motor domain construct is sufficient to crosslink and sort MTs into parallel bundles *in vitro*, which are general features of an MT organizer. Electron microscopy (EM) and hydrodynamic analysis reveal that KLIF is a parallel dimer suggesting a novel motility mechanism to sort and organize microtubules. The KLIF motor domain contains a series of active site mutations found in other MT-organizing motors that slow the overall rate of the motor but increase its bundling capacity. An intrinsically disordered N-terminal extension serves as a secondary MT binding domain and is essential for bundling. In addition to crosslinking MTs, KLIF forms stable links between MT plus ends, thereby aligning their ends. Together, these results indicate that KLIF may function in the cell to form a bundle of aligned MT plus ends that make up the new posterior end just prior to the completion of cytokinesis. These studies show at a mechanistic level how the activities of a novel MT organizing kinesin is integral to the completion of division and transmission of cell shape.

## Results

### KLIF is a double-headed non-processive motor

The domain structure of KLIF is similar to most N-terminal kinesins, which serve diverse functions including cargo transport and MT organization (16). KLIF contains an N-terminal catalytic motor domain that binds MgATP and MTs. The motor domain also contains an intrinsically disordered region of 126 amino acids at its N-terminus (NT) that is enriched in proline, serine, threonine, and basic residues. The motor domain is followed by a stalk that is composed of ∼800 amino acids of predicted coiled-coil sequence, which generally serves to oligomerize the heavy chains of other kinesin classes to ensure that pairs of motors can take coordinated steps on MT tracks (17). The second half of the stalk domain is composed of amino acid repeats which are followed by a small globular tail at the C-terminus that lacks homology to any known domain (**Fig. 1A**).

**Figure 1.**
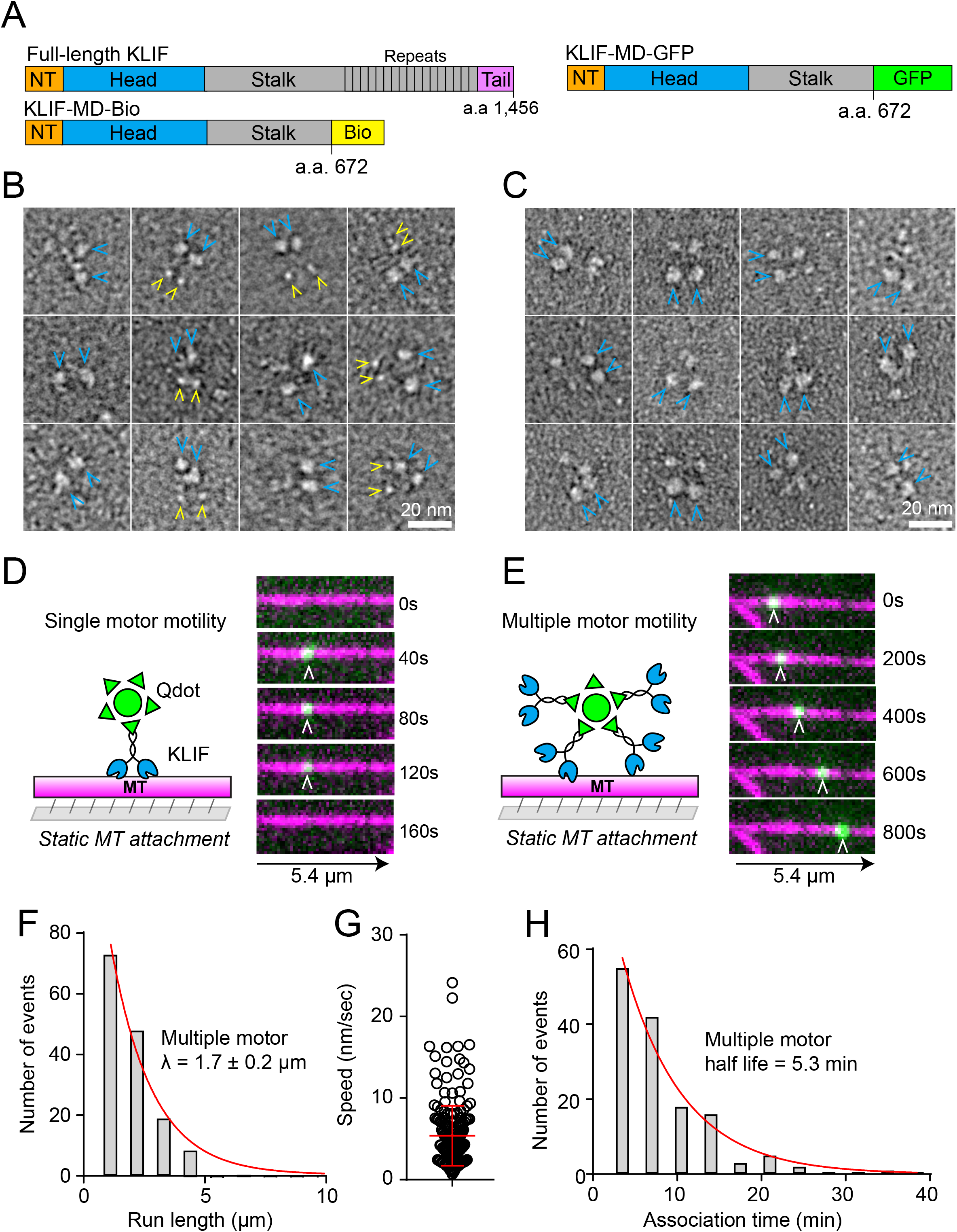
Negatively stained EM images of KLIF motor domain constructs and experimental setup for single and multiple motor motility assays of KLIF on microtubules. (A) Domain structure of full-length KLIF compared to motor domain constructs fused to C-terminal biotin tag (KLIF-MD-Bio) or GFP tag (KLIF-MD-GFP). Gallery of representative images of (B) KLIF-MD-Bio and (C) KLIF-MD-GFP negatively stained and imaged by EM. Blue arrowheads indicate the KLIF motor domain, which are typically observed in pairs. Pairs of smaller globular structures (yellow arrowheads) are observed in a subset of KLIF-MD-Bio images. Experimental setup and TIRF microscopy images showing (D) no movement of a single KLIF-MD-Bio bound to a Qdot (green) on MTs (magenta) versus (E) continuous movement of small motor ensembles bound to a Qdot at 1 mM MgATP. (F) Frequency distribution of run lengths for KLIF-MD-Bio motor ensembles moving on MTs at 1 mM MgATP. Characteristic run length, λ, was determined by fitting the distribution to an exponential function (y = Ae^−x/λ^) (red line). Error for λ is in SE of the fit. (G) Speed distribution of KLIF-MD-Bio motor ensembles bound to a Qdot. The average speed is a geometrical average to a Gaussian fit to the binned data is 5.4 ± 3.7 nm/sec SD. (H) Distribution of motor association times with MTs. Half-life was determined using a fit to an exponential curve Y=Ae^-k*x^.

To examine the properties of KLIF *in vitro*, we expressed and purified a construct containing the KLIF motor domain and a ∼170 amino acid portion of the stalk fused to a C-terminal biotin (KLIF-MD-Bio) or GFP tag (KLIF-MD-GFP) (**Fig. 1A, Supplemental figure S1**). These constructs lack the globular tail and repetitive sequences in the stalk in order to simplify our analysis of the motor. We first used negative stain electron microscopy (EM) to determine the stoichiometry and organization of KLIF-MD-Bio and KLIF-MD-GFP constructs. Images of KLIF-MD-Bio showed two distinct globular densities at one terminus (**Fig. 1B, blue arrowheads, Supplemental figure S2A**), consistent with a two-headed dimer. In a subset of these images, smaller nearby structures (**Fig. 1B, yellow arrowheads**) could be observed, which are likely the C-terminal biotin tags. Pairs of heads were also observed with KLIF-MD-GFP (**Fig. 1C, blue arrowheads, Supplemental figure S2B**). Because the two motor domains are together on one terminus, these data indicate that the KLIF is likely a parallel dimer.

We explored the possibility that KLIF-MD-Bio dimers could further oligomerize in solution at higher protein concentrations than what is used for EM analysis. To test this possibility, we performed sedimentation velocity experiments using a range of protein concentrations (**Supplemental figure S3A-B**). Under all concentrations tested, KLIF MD-Bio sedimented as a major species with an S-value of ∼6.0-6.3S, which corresponds to a molecular weight of approximately 150-185 kDa, indicating that KLIF-MD-Bio is most likely a dimer in solution. The sample exhibited a high frictional coefficient (f/f_0_=1.64-1.79) with a broad major peak, which indicates an extended conformation, consistent with the EM structures (**Fig. 1B-C**). These data show that there is no concentration dependence on the S-value up to 48.7 μM (**Supplemental figure S3B**).

We next visualized the movement of KLIF on MTs by conjugating the KLIF-MD-Bio construct to streptavidin quantum dots (Qdots) (**Fig. 1D-E**). These nanometer-sized fluorescent semiconductor crystals function as an artificial cargo and allow the formation of complexes containing one or more motors which can be visualized by fluorescence microscopy. Total internal reflection fluorescence (TIRF) microscopy was used to track the movement of Qdot-bound motors along rhodamine-labeled MTs immobilized inside a flow chamber. Cargo transporting motors are typically processive, which allows them to walk micron-long distances on cytoskeletal tracks as a single molecule without dissociating. To determine if KLIF can move processively on MTs, the motor was conjugated to streptavidin-coated Qdots through its C-terminal biotin tag at a ratio of 1 motor to 5 Qdots to ensure that the majority of Qdots are bound to a single motor (**Fig. 1D**). Under these conditions we observed transient associations of Qdots with MT tracks but no movement (**Fig. 1D, Supplemental movie S1**), indicating that the motor is non-processive under the conditions of our assay. We next tested if teams of motors bound to a single Qdot can move continuously on MTs. KLIF-MD-Bio was mixed with Qdots at a 50:1 molar ratio to ensure saturation of the Qdot (**Fig. 1E**). Considering its occupancy and size, this allows each Qdot to recruit 4-6 motors (18). Small teams of KLIF support robust motion on MTs (**Fig. 1E, Supplemental movie S2**) with a characteristic run length of 1.7 ± 0.2 μm (**Fig. 1F**) and average velocity of 4.1 ± 2.2 nm/sec (**Fig. 1G**). Given the slow speed of the motor, the observed long run lengths require the motor ensembles to remain attached to the MT for long periods of time. Fitting association time distributions to an exponential decay shows that motors stay associated with MTs while moving with a half-life of over 5 minutes (**Fig. 1H)**.

### KLIF contains features that optimize it for MT crosslinking

Because KLIF is slow compared to other cargo-transporting kinesins (which typically support speeds of ∼500 nm/sec) and not able to carry out processive transport on a single MT, it is unlikely that this motor traffics cargo in *T. brucei*. Instead, we considered that KLIF may function to crosslink MT networks in a manner analogous to mitotic kinesins. To test this hypothesis, the KLIF motor domain construct fused to GFP (KLIF-MD-GFP) and the cargo-transporting kinesin I motor domain construct fused to GFP (Kinesin I-MD-GFP) were purified and used in an *in vitro* MT bundling assay (**Fig. 2A-B, Supplemental figure S1**). The motors were premixed with rhodamine-labeled MTs in the presence of MgATP, adhered to flow chambers and the resulting MT structures were imaged using epifluorescence microscopy. KLIF has a strong tendency to bundle MTs (**Fig. 2B**). In contrast, bundling was not observed in the absence of motor or with Kinesin I-GFP under the same conditions employed for KLIF (**Fig. 2B**). Taken together, these results indicate that KLIF has the properties of a MT organizer rather than a cargo transporter.

**Figure 2.**
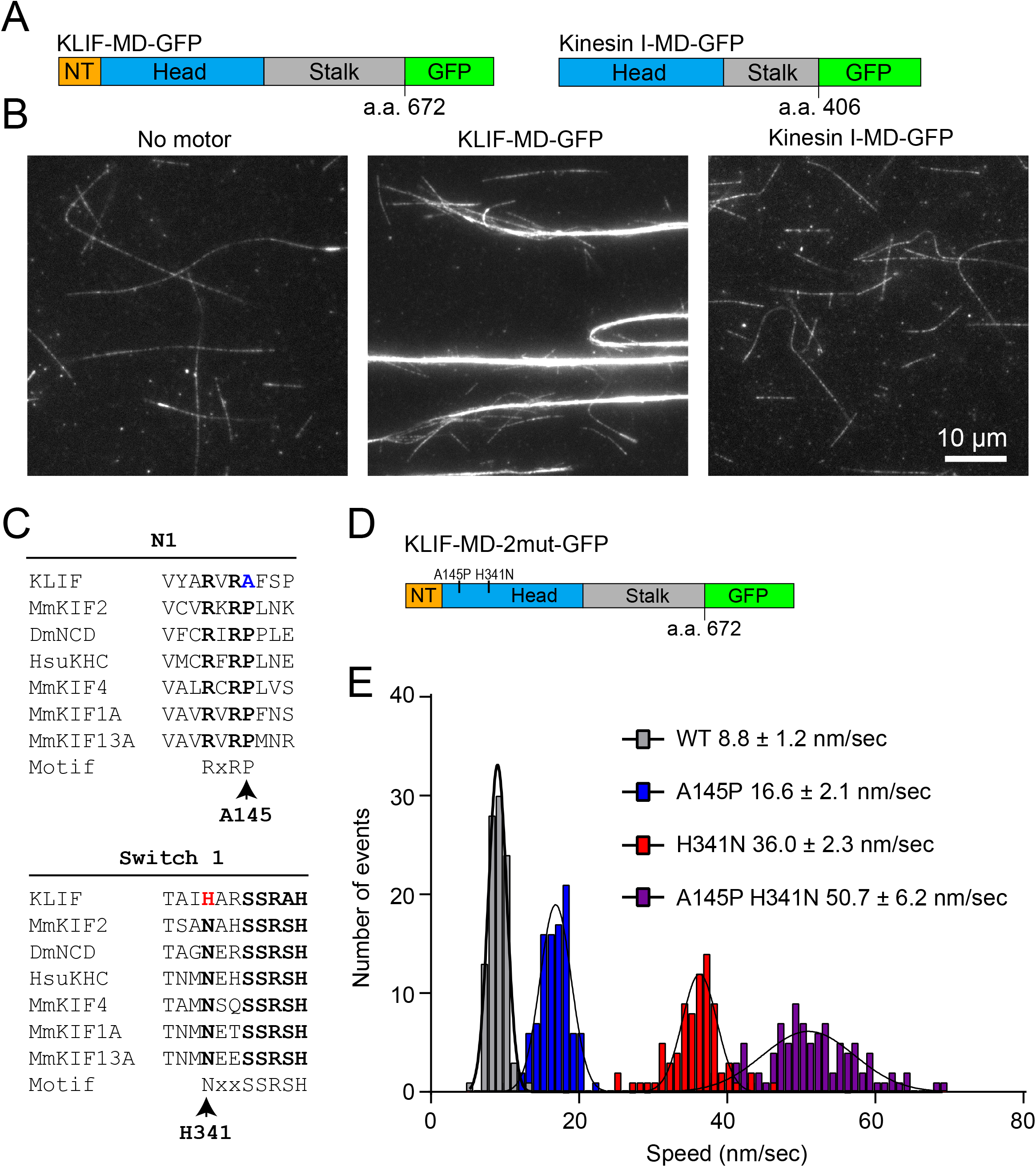
Images of MTs in the presence of KLIF or Kinesin I and speed distributions of expressed KLIF motor domain constructs carrying point mutations. (A) Schematic of the domain structure of the KLIF motor domain (KLIF-MD-GFP) and the Kinesin I motor domain (Kinesin I-MD-GFP) containing a C-terminal GFP tag. (B) Epifluorescence images showing rhodamine-labeled MTs without motor, MTs with 150 nM KLIF-MD-GFP, or MTs with Kinesin I-MD-GFP, mixed an incubated for 20 minutes in the presence of 1 mM MgATP, and applied to a flow chamber for imaging. (C) Alignment of KLIF N1 and switch I motifs showing divergence in conserved residues with other kinesin classes. (D) Schematic of the domain structure of a KLIF-MD-GFP containing A145P and H341N substitutions (KLIF-MD-2mut-GFP). (E) Representative speed distributions of WT (n=100) and mutant KLIF-MD-GFP motor domain constructs containing the A145P (n=101), H341N (n=79) or both (n=98) point mutations in an *in vitro* motility assay. Mean ± SD is shown. Means are significantly different from WT (P < 0.0001, t test comparing the Gaussian distributions).

KLIF has a set of amino acid substitutions in the catalytic domain that are invariant across a broad set of kinesins from other organisms. A sequence alignment with other well-characterized kinesins shows that KLIF lacks a well-conserved proline in the adenine base-binding N1 motif (RxRP). Other kinesins that lack this conserved proline include several BimC family members that function in spindle maintenance. This class of motors crosslinks MTs and are characteristically slow (19-21). In addition to this site, KLIF also contains an asparagine to histidine substitution in the switch I motif that is involved in hydrolysis of the γ-phosphate bond of MgATP (22, 23) (**Fig 2C**). We sought to determine if these unusual mutations could be contributing to slow rate and MT-organizing properties of the KLIF motor domain.

KLIF-MD-GFP domains containing point mutations in the RxRP motif (A145P) and switch I (H341N), which restore these amino acids to their consensus motifs, were expressed and purified to understand how these substitutions tune the activity of the motor (**Fig. 2D, Supplemental figure S1**). An *in vitro* gliding filament motility assay was used to determine if these changes altered the speed of the motor. In this assay, the motor is attached to the surface of a flow chamber via the GFP to orient the heads away from the coverslip. Rhodamine-labeled MTs are added to the chamber and are captured by KLIF. After washing away unbound MTs, motility buffer containing 1 mM MgATP was added to activate the motor. Filament gliding across the surface over time was monitored by epifluorescence microscopy. We found that the A145P mutation resulted in a ∼2-fold increase in speed (16.6 ± 2.1 nm/sec) compared to WT (8.8 ± 1.2 nm/sec), while the H341N mutation results in a ∼4-fold increase in velocity (36.0 ± 2.3 nm/sec). Mutating both residues (A145P, H341N) results in a ∼5-6-fold increase in velocity (50.7 ± 6.2 nm/sec), suggesting that the mutations result in an additive increase in rate (**Fig. 2E**). It is worth noting that while this enhanced speed is significantly faster than wild type KLIF, it is still ∼10-fold slower than conventional cargo-transporting kinesins (24). We identified two additional amino acids in switch I (A347) and switch II (C379) that diverge from conserved sequence motifs. However, mutating these to residues had minimal effects on the speed of the motor (**Supplemental figure S4**).

Along with unique mutations within the motor domain, KLIF has an intrinsically disordered N-terminal extension (NT) that is rich in proline and basic residues (**Fig. 3A**). Disordered N-terminal extensions are present in several of the kinesin-5 family of motor proteins, which function in bipolar spindle assembly and elongation, and appear to enhance their MT binding properties, which is likely an important feature for maintaining MT crosslinks (25, 26). The N-terminal sequence fused to a GFP-Bio and 8x HIS tag (KLIF-NT-GFP-Bio) was expressed and purified to determine if the N-terminal extension of KLIF can bind MTs on its own. The same construct lacking the N-terminal KLIF sequence (GFP-Bio) was purified as a control (**Fig. 3A, Supplemental figure S1**). KLIF-NT-GFP-Bio or GFP-Bio were added to a blocked flow chamber with surface-adhered rhodamine-labeled MTs, followed by epifluorescence microscopy to monitor the GFP signal. The KLIF-NT-GFP-Bio bound to MTs, while minimal binding was seen with the GFP-Bio control. These data indicate that the N-terminal domain contains an MT-binding domain that is independent of the motor domain (**Fig. 3B**).

**Figure 3.**
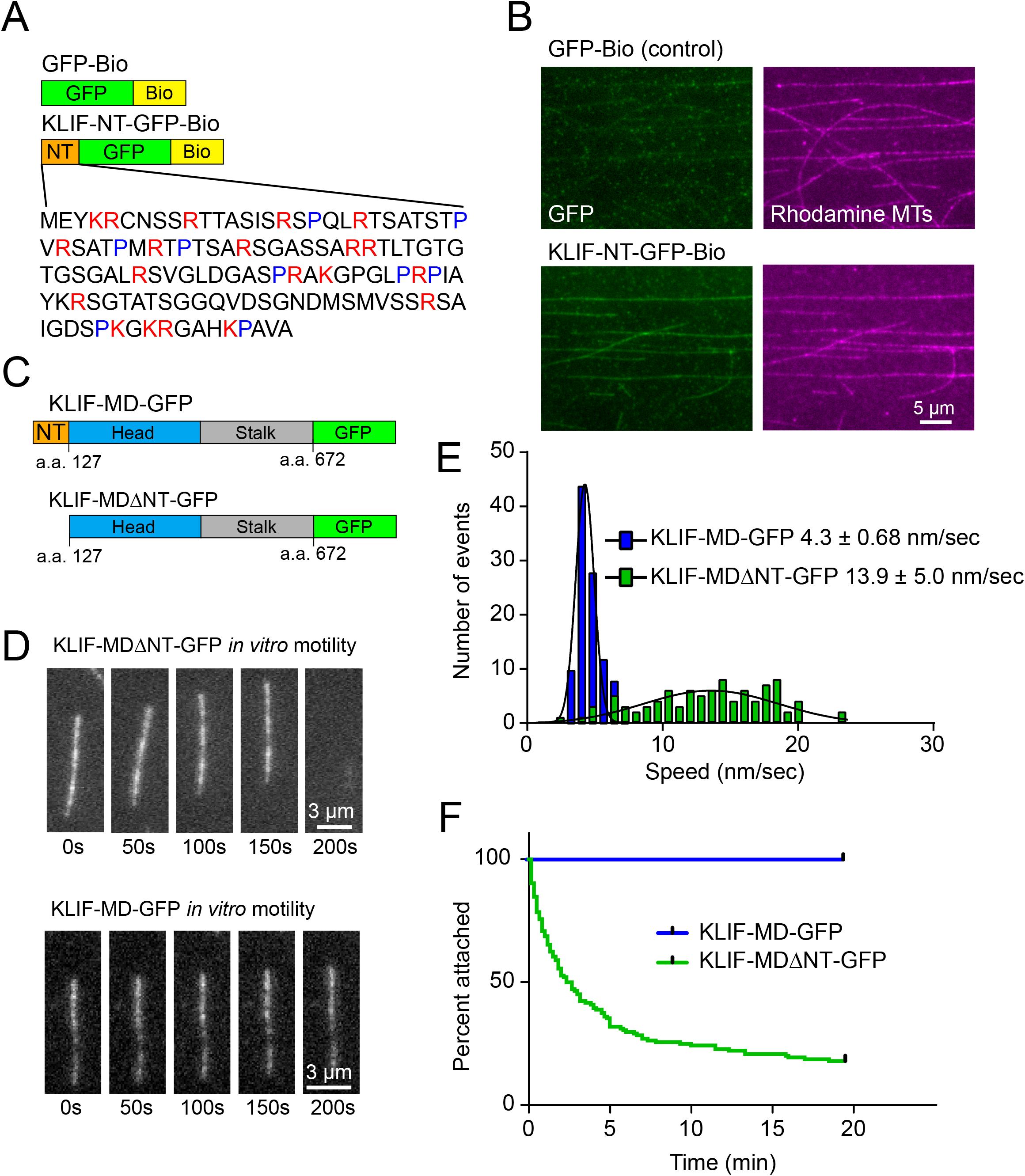
MT binding of the N-terminal extension of KLIF and motility of a truncated KLIF construct lacking its N-terminal extension. (A) Domain structure of GFP fused to biotin (GFP-Bio) and the 126 amino acid N-terminal extension of KLIF fused to GFP-biotin (KLIF-NT-GFP-Bio). The sequence of the N-terminus is shown with proline residues indicated in blue and basic amino acids in red. (B) Epifluorescence microscopy images showing that the N-terminal extension of KLIF fused to GFP-biotin (KLIF-NT-GFP-Bio) but not GFP-biotin (GFP-Bio) (left panel) interacts with rhodamine-labeled MTs *in vitro* (right panels). (C) Schematic of the domain structure of a KLIF construct lacking its N-terminal extension (KLIF-MDΔNT-GFP). (D) Epifluorescence images showing rhodamine-labeled MT gliding by (top) KLIF-MDΔNT-GFP versus (bottom) KLIF-MD-GFP in an *in vitro* motility assay. (E) Speed distribution comparing KLIF-MD-GFP (blue; n=102) and KLIF-MDΔNT-GFP (green; n=93) *in vitro* motility. Mean ± SD is shown. (F) Kaplan-Meier survival plot comparing MT dissociation times during an *in vitro* motility assay for KLIF-MD-GFP (blue; n=102) and KLIF-MDΔNT-GFP (green; n=144). Survival curves are significantly different (P<0.0001 using a Mantel-Cox test).

The presence of two MT binding domains at the N-terminus of KLIF suggests that they may work in tandem to enhance motor binding. To determine the contribution of the N-terminal extension, the KLIF motor domain lacking the N-terminal extension fused to GFP (KLIF-MDΔNT -GFP) (**Fig. 3C, Supplemental figure S1**) was expressed, purified, and then used in gliding filament motility assays. High motor densities were necessary for KLIF-MDΔNT-GFP to capture MTs to the surface of the imaging chamber. This construct supported gliding in the presence of MgATP with a broad speed distribution averaging 13.9 ± 5 nm/sec (**Fig 3D-E**). Under the same surface motor densities, KLIF-MD-GFP translocated filaments with an average speed of 4.3 ± 0.68 nm/sec (**Fig 3D-E**), which is reduced compared to previous *in vitro* motility assays (**Fig 2C**) due to the high motor densities used in this experiment. These data indicate that the motor can interact with MTs independent of its N-terminal extension and support gliding. However, in gliding assays using the KLIF-MDΔNT-GFP construct, MTs were not persistently attached compared to WT and dissociated from the surface over time (**Fig 3D, Supplemental movie S3-S4**). We measured dissociation times of MTs in the *in vitro* motility assay and plotted these times in a Kaplan-Meier survival plot. Over 50% of the filaments dissociated from surface bound KLIF-MDΔNT-GFP within 2.5 minutes, while no filaments dissociated from wild type motors under the same conditions (**Fig 3F**). This indicates that the N-terminal extension is not required for motility but functions to enhance MT binding.

The KLIF motor domain mutants (**Fig. 4A**) were tested for changes in their ability to bundle MTs. Different concentrations of KLIF constructs were incubated with rhodamine-biotin-labeled microtubules in solution for 20 minutes. The microtubules were then captured to the surface of a blocked biotin-neutravidin coated flow chamber and imaged using epifluorescence microscopy. In control experiments, KLIF-MD-GFP bundled MTs down to a concentration of 40 nM (**Fig. 4B-C, red, Supplemental figure S3**). The KLIF-MD-2mut-GFP mutant, which enhanced the *in vitro* motility speed of the motor ∼5-fold, showed a significantly reduced bundling activity at 150 nM and 40 nM compared to wild type (**Fig. 4B-C, blue, Supplemental figure S3**). The KLIF MD construct lacking the N-terminal extension (KLIF-MDΔNT-GFP) showed no MT bundling at any concentration, indicating that the N-terminal extension is required for KLIF bundling activity (**Fig. 4B-C, grey, Supplemental figure S3**). A construct containing just the KLIF N-terminal extension (KLIF-NT-GFP-Bio) did not bundle MTs at any protein concentration (**Fig. 4B-C, magenta, Supplemental figure S3**). From these data we can conclude that both the motor domain and its N-terminal extension are required for MT bundling *in vitro*, and that the mutations that enhance the speed of the motor diminish its ability to bundle.

**Figure 4.**
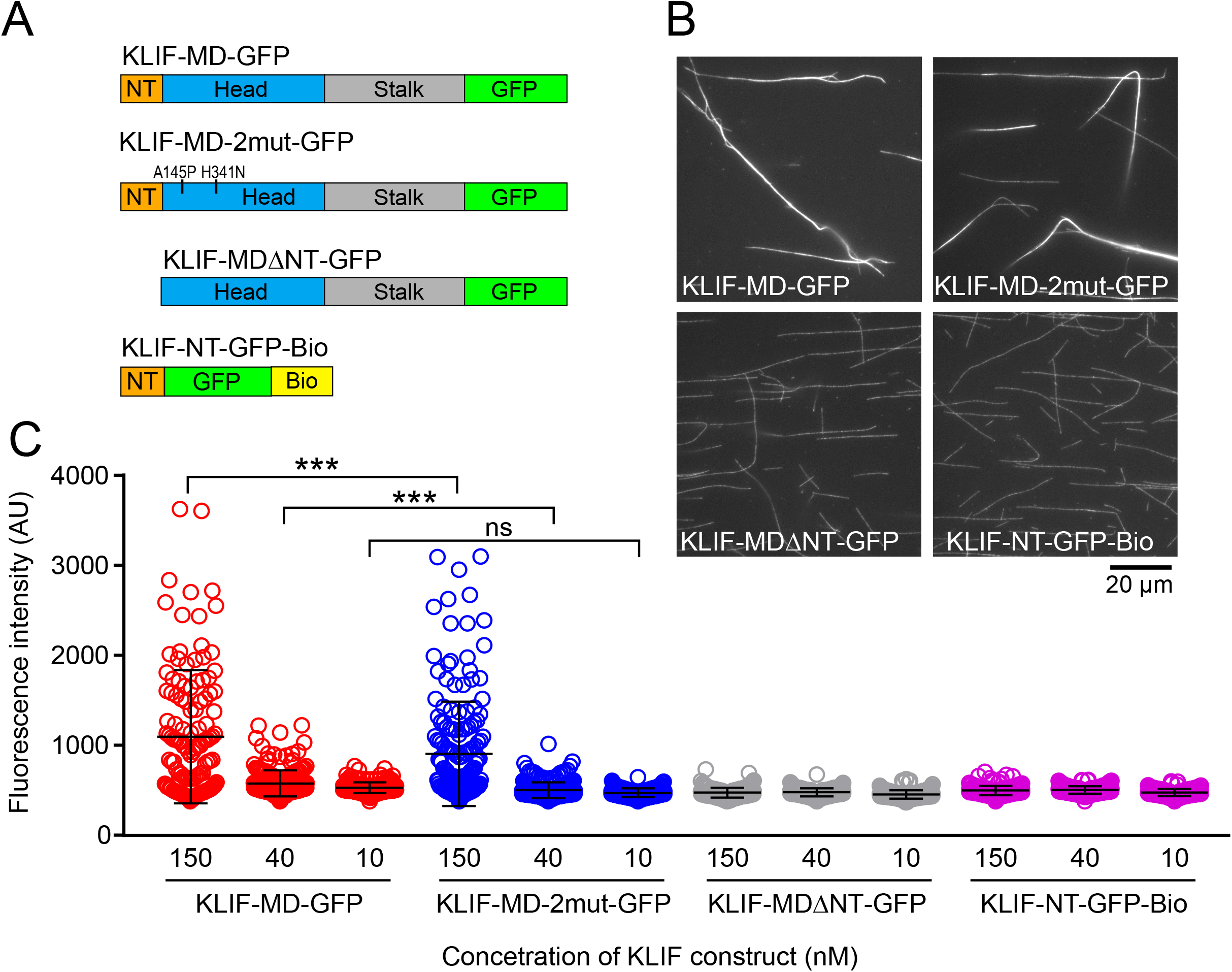
Epifluorescence imaging of MTs bundled by KLIF. (A) Domain structure of KLIF constructs used in MT bundling experiments. (B) Epifluorescence images showing the resulting MT organization when mixing rhodamine-labeled MTs with 150 nM of the indicated KLIF construct in solution for 20 minutes in the presence of 1 mM MgATP and applied to a flow chamber. (C) Scatter plot showing the fluorescence intensity distributions of rhodamine-labeled MTs mixed with KLIF constructs at 150 nM, 40 nM and 10 nM of the indicated KLIF construct. Intensity measurements were generated by taking maximum intensity pixel over a cross section of each MT structure. Mean intensity ± SD for KLIF MD-GFP (red) at 150 nM (1089 ± 741.2, n = 133), 40 nM (559.2 ± 144.4, n = 202), 10 nM (483.2 ± 59.4, n = 217). For KLIF MD 2mut-GFP (blue) at 150 nM (851.2 ± 581.1, n = 172), 40 nM (458.4 ± 86.9, n = 232), 10 nM (472.8 ± 48.9, n = 214). For KLIF MD ΔNT-GFP (green) at 150 nM (457.5 ± 55.48, n = 216), 40 nM (430.8 ± 44.6, n = 216), 10 nM (432 ± 45.5, n = 221). For KLIF NT-GFP-Bio (magenta) at 150 nM (460.9 ± 50.91, n = 225), 40 nM (413.5 ± 42.8, n = 218), 10 nM (434.3 ± 39.9, n = 219). The fluorescence intensity of MTs bundles with KLIF MD-GFP is statistically greater at 150 nM and 40 nM concentrations compared with KLIF MD 2mut-GFP (P < 0.05, Mann–Whitney test).

### KLIF sorts MTs into parallel bundles *in vitro*

MT crosslinking proteins can produce MT bundles with the MTs arranged in either a parallel or anti-parallel orientation. Each of these arrangements would provide important information about KLIF’s ability to organize MTs, so we devised a strategy to determine the MT orientation within bundles using a modified *in vitro* motility assay adapted from Braun *et al* (27). MTs were polarity marked at their minus-ends by growing rhodamine-labeled tubulin (**Fig. 5A, red**) from Cy5-labeled seeds (**Fig. 5A, green**). Bundles were first formed in solution by mixing KLIF-MD-Bio with MTs in the presence of MgATP for 40 minutes to provide sufficient time for sorting. The bundles were then added to a flow chamber and adhered to the surface by KLIF, forming a dynamic attachment. Unlike in previous assays which used a static attachment, surface bound KLIF is capable of dissociating bundles into individual filaments in the presence of MgATP using an *in vitro* motility assay (**Fig. 5A**). Because KLIF can only move MTs with their minus-ends leading (13), the orientation of each filament in the bundle is determined by the direction of their disassembly. We observed predominantly unidirectional disassembly (**Fig. 5B, Supplemental movie S5**) which was quantitated by measuring the angle of disassembly after 5 minutes relative to the axis of the bundle at time 0 (**Fig. 5B**). Using this approach, angles approximating 0° exit the bundle forwards and are parallel while angles near 180° result from filaments running anti-parallel in the bundle. To simplify the analysis, only bundles less than 30 μm in length were measured. Binning the angles on a polar coordinate plot shows that filaments dissociate from the bundle unidirectionally (**Fig. 5C**), which argues that KLIF forms bundles that are almost exclusively composed of parallel MTs.

**Figure 5.**
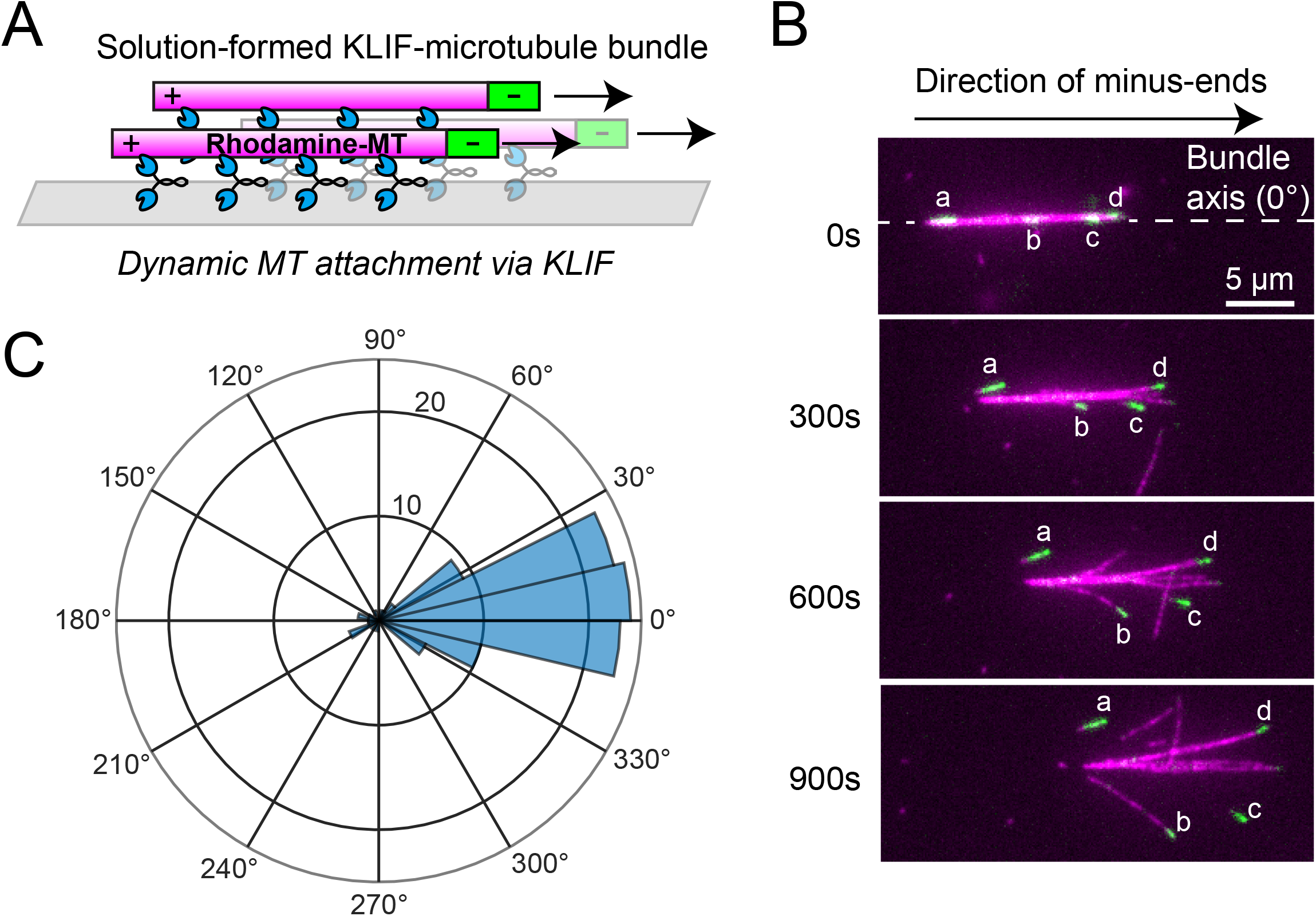
Modified *in vitro* motility assay schematic and polar coordinate plot showing angles of bundle disassembly. (A) Schematic of a modified *in vitro* motility assay to determine bundle polarity. Bundles are formed with polarity marked MTs in solution for 40 min in the presence of 1 mM MgATP and applied to a flow chamber. Once attached, the bundle disassembles by KLIF *in vitro* motility and the orientation of each filament is determined by the direction gliding relative to other filaments in the bundle. (B) Epifluorescence microscopy images showing a bundle formed by incubating polarity marked (green, letters) MTs, unidirectionally disassembling in an *in vitro* motility assay in the presence of 1 mM MgATP. Unidirectional disassembly indicates that the bundle is formed from parallel MTs. (C) Polar coordinate plot showing the angles of disassembly from the bundle after 5 minutes of *in vitro* motility (n = 109). The axis was determined after initial attachment of the bundle to the surface of a flow chamber. Angles approximating 0° indicate filaments that were parallel in the bundle and angles approximating 180° indicate filaments that were oriented antiparallel.

To probe the mechanism KLIF employs to construct parallel MT bundles, we used an assay that visualizes the dynamics of a simplified bundle comprising two MTs that are crosslinked in a parallel (**Fig. 6A**) or antiparallel (**Fig. 6D**) orientation. Polarity marked MTs containing biotin were attached to a neutravidin-bound flow chamber and washed to clear unbound MTs. KLIF-MD-GFP and polarity marked MTs without biotin were then added at concentrations that promote the crosslinking of non-biotinylated MTs with surface-adhered MTs. We then imaged the dynamics of crosslinked MTs using epifluorescence microscopy in the presence of 1 mM MgATP. When MTs are in a parallel orientation, they remained static over long periods of imaging (**Fig. 6B**). The two-MT overlap that ends at a polarity marked minus-end can be clearly observed when taking an intensity line scan over the MT pair (**Fig. 6C**). When MTs were crosslinked in an antiparallel orientation (**Fig 6D**), KLIF translocated the filaments relative to each other (**Fig. 6E, Supplemental movie S6**) with an average speed that is approximately double relative to single filament gliding in an *in vitro* motility assay (**Fig. 6F**). This doubling of speed indicates that KLIF walks on both of the MTs that it crosslinks.

**Figure 6.**
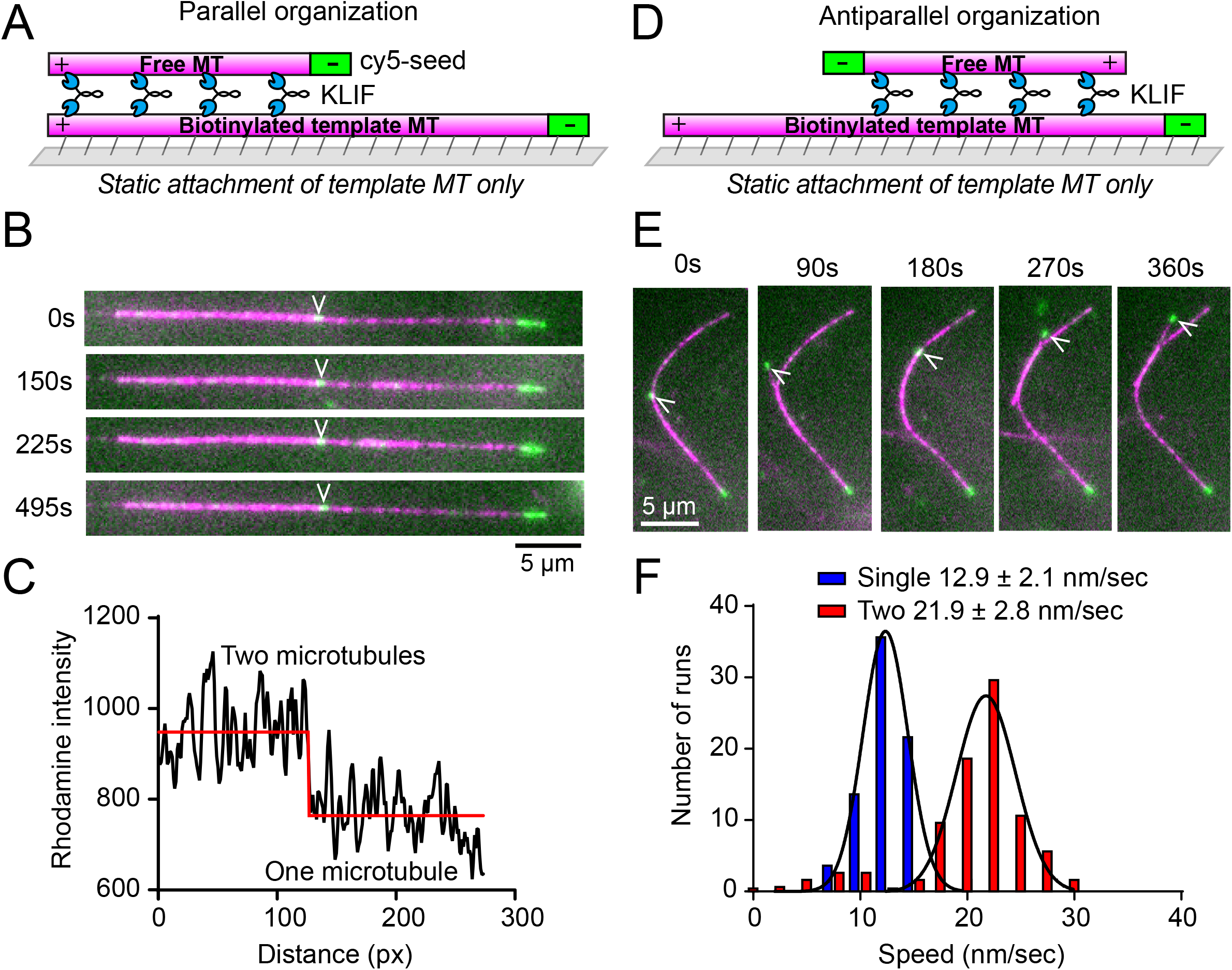
Two MT *in vitro* motility assay schematic and speed distributions for two MT movement compared to KLIF *in vitro* motility speeds. (A) Experimental setup for imaging the dynamics of parallel arranged MTs over time. (B) Epifluorescence microscopy images showing parallel arranged MTs (magenta) crosslinked by KLIF are non-motile. The polarity marked end (green) of the non-surface filament (free filament) is indicated with a white arrowhead. The template MT is biotinylated for attachment to the surface through biotin-streptavidin. (C) Intensity profile referring to the filament in B showing the location of two-MT overlap. The red line was generated using a custom step-finding algorithm. (D) Experimental setup for imaging the dynamics of antiparallel arranged MTs over time. (E) Epifluorescence microscopy images showing the movement of the free filament relative to the surface bound filament that is crosslinked by KLIF in an antiparallel orientation to a surface bound filament. (F) Speed distributions of KLIF MT gliding in a classic *in vitro* motility assay (blue) (n = 113) compared to the speeds of MT movement in a two-MT assay (red) (n = 89). Mean ± SD is shown.

### KLIF forms stable links between MT plus-ends

When antiparallel MTs gliding relative to each other aligned their plus ends in the two-MT assay, we frequently observed long-lived associations between MT plus ends that resulted in pivoting of the free filament about a fixed point with a high degree of rotational freedom. This is illustrated by the multiple orientations the free filament exhibits relative to the immobilized MT (**Fig. 7A, plus-end to plus end tethering, Supplemental movie S7**). On occasion, we also observed instances where a plus end associates with the minus end of an immobilized filament (**Fig. 7A, capture**). However, this interaction results in antiparallel overlap that will undergo relative gliding until the plus ends align (**Fig. 7A**). Plus end to plus end tethering typically results in dissociation of the filament or bundling in a static parallel orientation that aligns plus ends. Measuring the tethering times of multiple events and combining them into a histogram shows that these events are long lived with a fitted half-life of 1.7 minutes (**Fig. 7B**). These tethering times are considerably longer than the MgATP hydrolysis rate of KLIF (∼1 sec^-1^) (based on its *in vitro* motility speed) which suggests the existence of a static binding mode at the end of the filament that is distinct from interactions along the length of the MT that support gliding.

**Figure 7.**
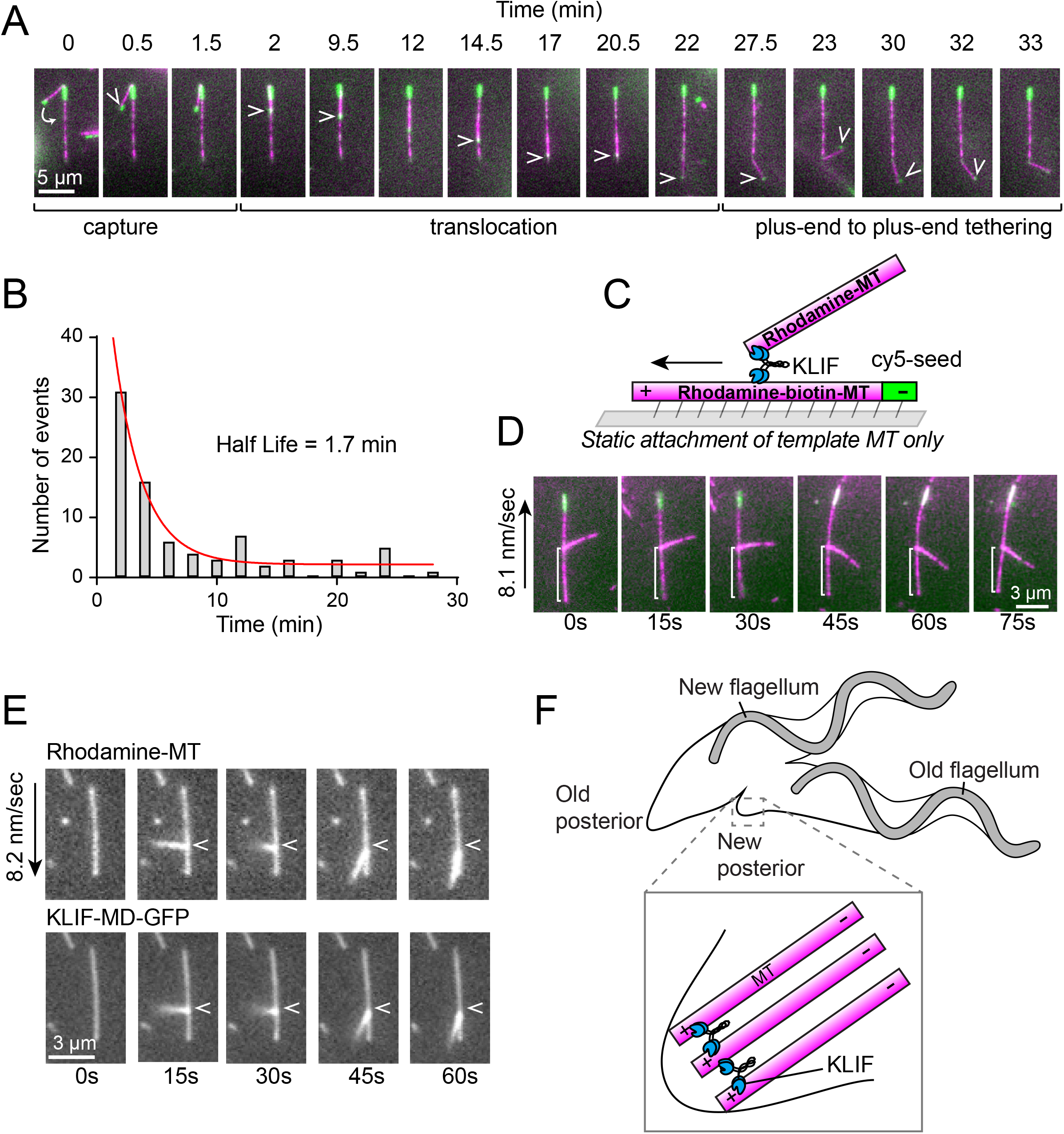
MT tethering and transport by KLIF and negatively stained EM images of KLIF. (A) Epifluorescence microscopy images showing the capture of a free MT by its plus end to the minus end of a surface-bound MT in a two-MT assay. The free MT then becomes crosslinked by KLIF in an antiparallel orientation and translocated to the end of the surface bound filament whereby the plus ends are aligned and tethered. (B) Distribution of times MTs remained tethered to immobilized filaments. Half-life was generated from the fit (red). n = 101. (C) Schematic of a MT being transported at its end by KLIF to the plus end of an immobilized filament. (D) Epifluorescence microscopy images showing a MT attached to the side of immobilized MT by its end being transported to the plus end of a polarity marked MT at the rate of 8.1 nm/sec. The decreasing length of the white bracket indicates movement over time. The image was rotated to account for movement of the bottom filament halfway through imaging. (E) Epifluorescence microscopy images showing rhodamine (top) and KLIF-MD-GFP (bottom) imaging of a MT being transported by KLIF on a surface attached MT by its end. White arrow indicates the start of the run. (F) Model for how KLIF (blue) functions to complete cytokinesis in *T. brucei* by focusing MT plus-ends (magenta) to the site of the new posterior (inset).

Interestingly, we occasionally observed free filament plus ends tethered to the side of template MTs (**Fig. 7C-E**), which were subsequently transported by KLIF towards the plus end of immobilized MTs at a speed that is consistent with the *in vitro* motility rate of KLIF ensembles walking on a single MT (**Fig. 7D, Supplemental movie S8**). Imaging KLIF-MD-GFP while a MT end is being transported shows the accumulation of KLIF at the point where the MTs connect (**Fig. 7E**), suggesting that multiple motors likely facilitate the association of MT ends with the interacting filament. This is consistent with our findings that KLIF is non-processive and cannot support movement along a microtubule as a single molecule (**Fig. 3D-E**).

## Discussion

Depletion of KLIF in *T. brucei* results in a severe cytokinesis defect where the daughter cells fail to construct a new posterior end, which is essential for the completion of cell division (13). To understand how KLIF functions in this process, we studied the properties of a simplified motor domain construct of KLIF *in vitro*. Using single-molecule techniques, we showed that KLIF is non-processive because it is unable to move micron-long distances as a single molecule. Because single molecule processivity is a hallmark of a cargo transporter, we considered that KLIF may function instead to organize MTs much like the mitotic kinesins. Consistent with this idea, we found that KLIF has a strong tendency to crosslink MTs and to form MT bundles in the presence of MgATP, which is a feature of a MT-organizing motor. KLIF can also sort MTs into parallel bundles and stabilize the alignment of MT plus ends. Because of these unique features, we propose that KLIF functions to complete cytokinesis by remodeling MTs to form a bundle of aligned plus ends that define the shape of the newly emerging posterior end (**Fig. 7F**). Once these MTs are organized by KLIF, other MT crosslinking proteins likely function to stabilize the new structure.

### How does KLIF gather MT plus-ends at the site of the new posterior?

We show that KLIF can form static links between MT plus ends that persist for several minutes (**Fig 7B**), and in some instances KLIF can link the plus ends of MTs to the side of a template MTs. Considering KLIF is non-processive, these interactions are likely stabilized by a complex containing multiple motors. Time-lapse imaging of MT plus ends bound to the side of a template MT showed that KLIF is capable of plus end directed transport of these MTs with a speed that corresponds to the gliding filament speed. This indicates that KLIF forms a static complex at MT plus ends that is capable of transport along the stationary MT. Negatively stained images of the *T. brucei* subpellicular array show that short MTs become intercalated in the array late in the cell cycle (28). These MTs only associate with long MTs in the array by their ends. Given that KLIF can support plus end directed transport by their ends, it is an intriguing possibility that these short MTs may be transported by KLIF along MTs in the array to the site of the new posterior. This is one potential mechanism for how KLIF could gather MT plus ends to the site of the new posterior during cytokinesis. Xmap215, a MT plus end binding protein, enriches at the site of the new posterior in the late stages of cytokinesis in *T. brucei* (11). This localization corresponds temporally with the arrival of KLIF to the new posterior end (13), providing further evidence for the role of the motor in concentrating MT plus ends at the site of the new posterior.

### Comparisons with other kinesins

Despite a lack of sequence homology with known classes of kinesins, KLIF has remarkably similar biophysical properties to kinesins that function in the creation and maintenance of the mitotic spindle. Kinesin-5 proteins, which include Eg5, are essential for the bipolar organization of the mitotic spindle. Their tetrameric bipolar head configuration allows these kinesins to crosslink and slide interpolar MTs relative to one another (29, 30). For both KLIF and Eg5, this sliding activity depends on the relative orientation of the crosslinked MTs, with antiparallel MTs gliding and parallel MTs statically crosslinked. The Eg5 gliding speed for two antiparallel MTs relative to one another is twice its *in vitro* motility velocity, indicating that the motor walks on both of the MTs it crosslinks, which is a functional consequence of its bipolar head configuration (31). The speed of KLIF in the two-MT assay is also twice the sliding filament velocity, implying that KLIF also walks on both MTs it crosslinks. However, our EM and sedimentation data show that KLIF is organized as a parallel dimer. Thus, the mechanism it employs to organize MTs must be different from that of Eg5. Minus end directed Kinesin-14 motors also can slide antiparallel MTs relative to one another. This class of kinesins bind one MT through its head and another through an MgATP-independent binding site in the tail and moves collectively toward the minus-end to move MTs relative to one another (27, 32-36). Because the KLIF motor domain construct in this study lacks its tail, its mechanism of action must also differ from this class.

### Features that enable KLIF to crosslink MTs

We identified several features in KLIF that promote MT crosslinking. The first is an intrinsically disordered N-terminal domain (NT) that is sufficient for binding MTs. KLIF lacking the NT requires high surface densities to capture MTs to the surface of a motility chamber compared to the wild type motor domain. MTs captured by the NT mutant quickly dissociate after adding a buffer containing MgATP, indicating that the association of KLIF with MTs is weak in the absence of the NT. While MTs were held transiently to the surface, this construct supported motility, indicating that this NT domain is not essential for the catalytic activity of the motor. These results suggest that the function of the NT is to provide an additional MT-interacting domain to enhance the affinity of the motor, which is likely a necessary adaptation that allows the dimeric motor to maintain crosslinks with two MTs as it steps. Disordered N-terminal extensions are observed in other kinesins (25). Like KLIF, the mammalian kinesin-3 Kif14 also contains an intrinsically disordered N-terminal domain. This kinesin localizes to mitotic spindle midbody and its depletion causes failed cytokinesis but the mechanism for how this motor functions in cytokinesis is unknown (37, 38). Recent work has shown that the Kif14 NT is sufficient for crosslinking MTs and also confers processivity to the dimeric motor, enabling it to move in crowded environments (26). While the KLIF NT does not provide KLIF with processive movement and is not able to crosslink MTs in the absence of the motor domain, enhanced MT binding may be a necessary feature to allow KLIF to remain attached to MTs in crowded environments such as the trypanosome subpellicular array, which is heavily crosslinked by MT associated proteins (MAPs) (39).

Phosphoproteomic data indicate that 12 amino acids in the NT of KLIF are phosphorylated (40). Given that the NT is necessary for crosslinking MTs, phosphorylation may serve as a key regulatory mechanism to ensure spatial and temporal control over its activity during cytokinesis. Phosphorylation would most likely limit the ability of the NT domain to bind MTs due to the increased negative charge, which would weaken the interaction with the acidic C-terminal tails present on alpha and beta tubulins. A similar mechanism that may reduce MT binding has been proposed for the chromokinesin Kid, which is phosphorylated on the second MT-binding region within the stalk (41).

The MT crosslinking activity of KLIF is also influenced by mutations in highly conserved amino acid residues in the N4 and switch I motifs that reduce the speed of the motor ∼6-fold but enhance the bundling activity of the motor. These amino acid changes are conserved in KLIF orthologs the related parasites *Leishmania* and *Trypanosoma cruzi*, and the ortholog present in the free-living kinetoplastid *Bodo saltans*. The switch I change in KLIF, which reduces the velocity of the motor 4-fold, is likely involved in nucleophilic attack of the γ-phosphate bond of MgATP (23, 42), while the missing proline in the N4 (RxRP) motif that reduces the speed of the motor 2-fold is likely caused by changes in adenine binding (43). Interestingly, five other kinesin-5 family members lack this conserved proline (*S. pombe* Cut7, *Aspergillus nidulans* BimC, *Arabidopsis thaliana* ATFC1a, and *S. cerevisiae* Kip1 and Cin8), which may indicate a convergent evolutionary adaptation to enhance the crosslinking activity of MT-reorganizing kinesins at the expense of the speed of the motor (21).

This study presents the first biophysical description of any molecular motor in *T. brucei*. The biophysical features of KLIF are similar to those of well-studied MT reorganizing kinesins, yet the activity of the motor is utilized very differently: to remodel subpellicular MTs to create new posterior within the confines of an existing MT array. It is possible that the properties of these motors have convergently evolved due to the similarities in the MT architecture they create. Thus, studies on vastly different systems serve to inform our general understanding of how MT organizing motors function in cell division and morphogenesis.

## Methods

### Kinesin expression and purification

*T. brucei* KLIF (Tb927.8.4950) amino acids 1-672 was amplified by PCR from *Trypanosoma brucei* Lister strain 427 genomic DNA and inserted into the bacterial expression vector pET30a containing either a C-terminal biotin tag (KLIF MD-Bio) or mClover3 variant of GFP (KLIF MD-GFP) (44) and His8 tag for affinity purification. A KLIF construct containing only the intrinsically disordered N-terminal extension (NT-IDD) (amino acids 1-129) was fused to mClover3-biotin tag followed by a His8 tag and inserted into pET30a. As a control, the mClover3-biotin-His8 tag was cloned into pET30a. Kinesin-1 (NCBI:NM_008449.2) containing the N-terminal 406 amino acids of mouse kinesin-1 (kinesin406) was amplified by PCR and subcloned into pET30a containing either a C-terminal biotin-His8 tag or mClover3-His8 tag. A kinesin406 bacterial expression construct in pET21a containing a G235A “rigor” point mutation was a kind gift from Dr. Kathy Trybus. This mutant binds tightly to MTs but does not support motility in the presence of MgATP and is used for static attachment of MTs to the flow cell surface. Kinesin constructs were purified as described previously (13) with the exception that the protein was further purified by ion exchange using a Mono Q™ 5/50 GL column in buffer Q (50 mM Imidazole pH 6.7, 50 mM NaCl, 3 mM MgCl_2_, 1 mM EGTA, 0.1 mM EDTA, 2 mM β-mercaptoethanol, 0.05 mM MgATP) and eluted using a 50 mM to 500 mM NaCl salt gradient. The KLIF NT-IDD construct was further purified by ion exchange in buffer QT (50 mM Imidazole pH 8.0, 50 mM NaCl, 3 mM MgCl_2_, 1 mM EGTA, 0.1 mM EDTA, 2 mM β-mercaptoethanol) and eluted using a 50 mM to 500 mM NaCl salt gradient. For EM and analytical ultracentrifugation, KLIF protein preparations were dialyzed against buffer E (10 mM K-PIPES pH 7.4, 300 mM KCl, 2 mM MgCl_2_, 0.1 mM EGTA) containing 1 mM DTT and further purified using a Superdex 200 10/300 GL gel filtration column.

### Tubulin preparation

Cycled bovine tubulin (PurSolutions) was thawed briefly in a 37°C water bath and clarified at 400,000 x g for 6 min at 2°C. The concentration was determined using Bradford reagent and diluted to 100 μM in ice cold BRB80 (80 mM PIPES, pH 7.0, 0.5 mM EGTA and 2 mM MgCl_2_) supplemented with 1 mM GTP. Labeled MTs were generated by mixing 4μL of 20 μM Cy5 (PurSolutions) or Rhodamine (Cytoskeleton, Denver, CO) labeled tubulin with 9 μL unlabeled MTs and transferred to a 37°C water bath for 20 min and stabilized by adding 10 μM paclitaxel (Cytoskeleton, Denver, CO). MTs were stored at room temperature. The preparation of polarity marked MTs was adapted from previous methods (45) except Rhodamine-labeled tubulin was polymerized from Cy5 labeled seeds which generated Rhodamine-labeled MTs containing Cy5-labeled minus ends. Biotin-labeled tubulin was also included for generating polarity marked biotin labeled template MT used in two-MT assays.

### Single and multiple motor motility assays

MT gliding filament assays were done as described previously (13). For quantum dot (Qdot) motility assays, KLIF MD-Bio was diluted into buffer M (10 mM Imidazole pH 7.4, 4 mM MgCl_2_, 1 mM EGTA, 300 mM KCl, 0.1 mM MgATP), clarified 400,000 x g for 15 min at 4°C and its concentration was determined by Bradford. For singe molecule experiments, 33 nM KLIF MD-Bio was mixed with 167 nM Qdots (1:5 ratio) in clarification buffer containing 4 mg/mL BSA. At this ratio, the majority of Qdots are bound to a single motor. For ensemble experiments, 900 nM KLIF-Bio was mixed with 20 nM Qdots (45:1 ratio) which promotes the binding of multiple motors per Qdot. For attaching Rhodamine-labeled MTs, kinesin406 G235A (“rigor” kinesin) was absorbed to the surface of a flow chamber and then blocked with blocking buffer (BRB80, 10mM DTT, 0.5% K-casein, 2 mg/mL K-casein). Rhodamine-labeled MTs were then added and the flow chamber was washed with buffer B (BRB80, 10 mM DTT) containing 10 μM taxol. The KLIF/Qdot mixture was then diluted ∼1:400 in Go buffer (BRB80, 10 mM DTT, 2 mg/mL K-casein, 0.5% Pluronic F127, oxygen scavenging system (3 mg/ml glucose, 0.1 mg/ml glucose oxidase, and 0.18 mg/ml catalase), 2 mM MgATP, 10 μM taxol), added to the flow chamber and imaged every 2 seconds for 40 minutes using Total Internal Reflection Fluorescence (TIRF) microscopy.

### KLIF N-terminal extension MT binding assay

The KLIF N-terminal extension containing a C-terminal mClover3-biotin (KLIF NT-GFP-Bio) or mClover3-biotin (GFP-Bio) diluted into BRB80 containing 10 mM DTT and clarified at 400,000 x g for 20 min at 4°C. The concentration of clarified protein was determined using Bradford and diluted to 1 μM in ice cold imaging buffer (BRB80, 10 mM DTT, 2 mg/mL K-casein, 0.5% Pluronic F127, oxygen scavenging system (3 mg/ml glucose, 0.1 mg/ml glucose oxidase, and 0.18 mg/ml catalase), 10 μM taxol). For attachment of Rhodamine labeled MTs, kinesin406 G235A (“rigor” kinesin) was absorbed to the surface of a flow chamber and then blocked with blocking buffer (BRB80, 10 mM DTT, 2 mg/mL K-casein, 0.5% Pluronic F127). Rhodamine labeled MTs were added to bind the surface in BRB80 containing 2 mg/mL K-casein and then washed to remove unbound MTs. KLIF NT-GFP-Bio or GFP-Bio (control) was diluted 1/10 in imaging buffer and allowed to bind the MTs. The flow chamber was washed with imaging buffer to washout unbound protein. Rhodamine labeled MTs and mClover3 fusion proteins were imaged using epifluorescence microscopy.

### MT bundling and bundle dissociation assays

For MT bundling assays, KLIF containing a C-terminal mClover3 tag (KLIF MD-GFP) was clarified 400,000 x g for 15 min at 4°C and mixed at the indicated concentrations with 0.4 μM Rhodamine-biotin-labeled MTs in Go buffer (BRB80, 10 mM DTT, 2 mg/mL K-casein, 0.5% Pluronic F127, oxygen scavenging system (3 mg/ml glucose, 0.1 mg/ml glucose oxidase, and 0.18 mg/ml catalase), 2 mM MgATP, 10 μM taxol) for 20 minutes at room temperature. During the incubation, Neutravidin flow cells were prepared by first binding 0.5 mg/mL Biotinylated BSA, followed by blocking with blocking buffer (BRB80, 10 mM DTT, 2 mg/mL K-casein, 0.5% Pluronic F127) and then 0.05mg/mL Neutravidin. The flow cell was washed extensively with buffer B (BRB80, 10 mM DTT) to remove unbound Neutravidin. The KLIF MD-GFP and Rhodamine-biotin-labeled MT mixture was applied to a flow chamber for 3 minutes to allow MT structures to bind to the Neutravidin surface, washed 2 times with Go buffer and immediately imaged using epifluorescence microscopy.

Bundle dissociation assays were described previously (27). KLIF MD-Bio was clarified 400,000 x g for 15 min at 4°C and diluted to 150 nM with 0.4 μM polarity marked MTs and incubated 30 minutes at room temperature in Go buffer to form bundles in the presence of 2 mM MgATP. The mixture was added to an unblocked flow chamber to allow KLIF MD-Bio and bundled MTs to bind to the surface. The flow chamber was then washed with Go buffer to remove unbound motor and MTs. Addition of Go buffer containing MgATP allows the surface bound motors to disassemble MT bundles into individual MTs on the surface with their minus ends leading. Disassembly of bundles was imaged using epifluorescence microscopy every 10 seconds for 30 minutes.

### Two MT assay

Template polarity marked MTs containing biotinylated tubulin were bound to a Neutravidin coated flow chamber (described above) in the presence of 0.5 mg/mL K-casein and washed to remove unbound MTs. KLIF MD-GFP was clarified 400,000 x g for 15 min at 4°C and diluted to 90 nM with 60 nM polarity marked free MTs that do not contain biotinylated tubulin and immediately added to the flow chamber to promote crosslinking of free MTs to template MTs by KLIF MD-GFP. The flow chamber was then imaged using epifluorescence every 15 seconds for 30 minutes.

### Analytical ultracentrifugation

Gel purified KLIF MD-Bio was dialyzed overnight in buffer U (10 mM K-PIPES pH 7.4, 200 mM KCl, 2 mM MgCl2, 0.1 mM EGTA) at 4°C. Sedimentation velocity analysis was conducted at 20°C and 35,000 RPM using absorbance optics with a Beckman-Coulter Optima AUC analytical ultracentrifuge. Double sector cells equipped with quartz windows were used. The rotor was equilibrated under vacuum at 20°C and after an equilibration period of ∼30 minutes the rotor was accelerated to 35,000 RPM. Absorbance scans at 280 nm were acquired at 20 seconds intervals for ∼8 hours.

### Negative stain electron microscopy

Gel purified KLIF MD-Bio in buffer E (10 mM PIPES pH 7.4, 300 mM KCl, 2 mM MgCl_2_, 0.1 mM EGTA, 0.05 mM ATP, 1 mM TCEP) was diluted to 10–25 nM in BRB80 containing 1 mM TCEP. The diluted sample was applied to UV-treated, carbon-coated copper grids and stained with 1% uranyl acetate. Micrographs were recorded using an AMT XR-60 CCD camera at room temperature on a JEOL 1200EX II microscope at a nominal magnification of 60,000.

### Imaging and data analysis

Images were taken on a Zeiss Axio Observer.Z1 microscope (Carl Zeiss Microscopy— Oberkochen, Germany) equipped with epifluorescence and a spinning illumination ring VectorTIRF system (Intelligent Imaging Innovations, Inc.) housing 405/488/561/640 nm lasers. Both modalities use an Alpha Plan-Apochromat 100X/1.46 NA oil TIRF objective, Prime 95B Back Illuminated Scientific CMOS camera with a resolution of 110 nm/pixel (Teledyne Photometrics— Tuscon, AZ) and a Definite Focus.2 system (Carl Zeiss Microscopy) for automatic focus correction and run by Slidebook software (Intelligent Imaging Innovations, Inc.).

MTs and Qdots were tracked with ImageJ (National Institutes of Health – Bethesda, MD) using the particle-tracking plug-in MTrackJ (46). Intensity measurements of MTs were made using the straight-line feature in ImageJ. All frequency distributions were created in GraphPad (GraphPad software, LLC). Polar histograms were generated in using the Rose Plot function in MatLab (The MathWorks, Inc).

Analytical ultracentrifuge analysis was conducted using Sedfit, version 16.36. The direct boundary modeling program was used to fit individual data sets based on numerical solutions to the Lamm equation (47). The continuous sedimentation coefficient (c(s)) distribution plots were sharpened, relative to other analysis methods, because the broadening effects of diffusion are removed by use of an average value for the frictional coefficient. The c(s) analyses were done at a resolution of 0.05 S, using maximum entropy regularization with a 68% confidence limit.

## Acknowledgements

We thank Dr. Aoife Heaslip for her assistance with protein preparations and useful discussions. We thank Dr. Kathleen Trybus for the kinesin-1 bacterial expression vector. We thank Dr. Kenneth Campellone for assistance with gel filtration chromatography and Dr. Laurent Brossay for assistance with ion exchange chromatography. We thank the National Heart, Lung, and Blood Institute Electron Microscopy Core Facility and the Biophysics Core facility at the University of Connecticut’s Center for Open Research Resource & Equipment for the use of their facilities. We would like to thank Dr. Lillian Fritz-Laylin’s group for critical reading of this manuscript. This work was funded by the National Institutes of Health grants AI112953 and R56AI153336 to C.L.d.G.

---

Supplementary figure S1. **KLIF constructs used in this study**. SDS-PAGE gel showing the indicated KLIF construct used for *in vitro* assays. Asterisk denotes a GFP-containing degradation product that was confirmed using anti-GFP western blotting.

Supplementary figure S2. **A field of negatively stained images of (A) KLIF-MD-Bio and (B) KLIF-MD-GFP**.

Supplementary figure S3. **Sedimentation of KLIF-MD-Bio**. (A) Sedimentation plots showing that the indicated concentrations of KLIF MD-Bio sedimented with an S value of ∼6.0-6.3S which corresponds to a molecular weight of approximately 150-185 kDa. c(s) distributions were normalized to the max c(s) peak for the four samples. (B) Table showing peak type and percentage for the main peak seen for each sample concentration, as determined by the c(s) analysis.

Supplementary figure S4. **Speed distribution of mutant KLIF constructs**. (A) Alignment of KLIF switch I and switch II motifs with other kinesin classes. Speed distributions of KLIF-MD-GFP (n=100) compared to constructs containing a (B) A347S mutation (n=68) and (C) C379S mutation (n=101). Mean ± SD is shown. Means are significantly different from WT (P < 0.0001, t test comparing the Gaussian distributions).

Supplementary figure S5. **Epifluorescence imaging of MTs bundled by KLIF constructs at different concentrations of motor**. (A) Epifluorescence images showing the resulting MT organization when mixing rhodamine-labeled MTs with the indicated KLIF construct in solution for 20 minutes in the presence of 1 mM MgATP over a range of concentrations and applied to a flow chamber. (B) Scatter plot showing the fluorescence intensity distributions of rhodamine-labeled MTs mixed with KLIF constructs at 150 nM, 100 nM, 40 nM, 20 nM and 10 nM of the indicated KLIF construct. Intensity measurements were generated by taking maximum intensity pixel over a cross section of each MT structure. Mean intensity ± SD for KLIF-MD-GFP (red) at 150 nM (1089 ± 741.2, n = 133), 100 nM (1224 ± 893.4, n = 175), 40 nM (559.2 ± 144.4, n = 202), 20 nM (445.3 ± 47.5, n = 200), 10 nM (483.2 ± 59.4, n = 217). For KLIF-MD-2mut-GFP (blue) at 150 nM (851.2 ± 581.1, n = 172), 100 nM (781.3 ± 465.2, n = 216), 40 nM (458.4 ± 86.9, n = 232), 20 nM (448.5 ± 52.4, n = 228), 10 nM (472.8 ± 48.9, n = 214). For KLIF-MDΔNT-GFP (green) at 150 nM (457.5 ± 55.48, n = 216), 100 nM (466.4 ± 68.3, n = 213), 40 nM (430.8 ± 44.6, n = 216), 20 nM (428.7 ± 48.3, n = 222), 10 nM (432 ± 45.5, n = 221). For KLIF-NT-GFP-Bio (magenta) at 150 nM (460.9 ± 50.91, n = 225), 100 nM (431.9 ± 50.8, n = 217), 40 nM (413.5 ± 42.8, n = 218), 20 nM (400.3 ± 43.5, n = 216), 10 nM (434.3 ± 39.9, n = 219).

Supplementary movie S1. TIRF single molecule motility assay showing that KLIF-MD-Bio bound to Qdot (green) transiently associated with MTs (magenta) but does not support motility.

Supplementary movie S2. TIRF multiple motor motility assay showing that multiple KLIF-MD-Bio motors bound to a Qdot (green) support continuous movement on MTs (magenta).

Supplementary movie S3. *In vitro* motility assay with KLIF-MDΔNT-GFP showing that surface bound KLIF-MDΔNT-GFP supports MT gliding but MTs dissociate over time.

Supplementary movie S4. *In vitro* motility assay with KLIF-MD-GFP showing that surface bound KLIF-MD-GFP supports robust MT gliding.

Supplementary movie S5. Microtubule sorting assay. MTs (magenta) were polarity marked at their minus end (green) and incubated with KLIF-MD-Bio in solution in the presence of MgATP. Resulting bundled MTs were then applied to a KLIF-bound flow cell to dissociate MTs to determine how they were organized in the bundle.

Supplementary movie S6. Two microtubule assay showing the relative gliding of a non-biotinylated (free) polarity marked MT by KLIF-MD-Bio that is bound in an antiparallel orientation relative to a surface bound (template) polarity marked template MT.

Supplementary movie S7. Two microtubule assay showing capture of a free polarity marked MT by its plus end. This is followed by antiparallel association and relative gliding by KLIF-MD-Bio, plus end to plus end tethering and dissociation.

Supplementary movie S8. Two microtubule assay showing the movement of a microtubule end relative to an interacting polarity marked MT.

## Notes

### Competing Interest Statement

The authors have declared no competing interest.

## References

1. Wang X, Fu Y, Beatty WL, Ma M, Brown A, Sibley LD, Zhang R. Cryo-EM structure of cortical microtubules from human parasite Toxoplasma gondii identifies their microtubule inner proteins. Nat Commun. 2021;12(1):3065. doi: 10.1038/s41467-021-23351-1. PubMed PMID: 34031406; PMCID: PMC8144581.

2. Schwartz CL, Heumann JM, Dawson SC, Hoenger A. A detailed, hierarchical study of Giardia lamblia’s ventral disc reveals novel microtubule-associated protein complexes. PLoS One. 2012;7(9):e43783. doi: 10.1371/journal.pone.0043783. PubMed PMID: 22984443; PMCID: PMC3439489.

3. Hu K. Organizational changes of the daughter basal complex during the parasite replication of Toxoplasma gondii. PLoS Pathog. 2008;4(1):e10. doi: 10.1371/journal.ppat.0040010. PubMed PMID: 18208326; PMCID: PMC2211554.

4. Tumova P, Kulda J, Nohynkova E. Cell division of Giardia intestinalis: assembly and disassembly of the adhesive disc, and the cytokinesis. Cell Motil Cytoskeleton. 2007;64(4):288–98. doi: 10.1002/cm.20183. PubMed PMID: 17205565.

5. Schroeder TE. Actin in dividing cells: contractile ring filaments bind heavy meromyosin. Proc Natl Acad Sci U S A. 1973;70(6):1688-92. PubMed PMID: 4578441; PMCID: PMC433573.

6. Fujiwara K, Pollard TD. Fluorescent antibody localization of myosin in the cytoplasm, cleavage furrow, and mitotic spindle of human cells. J Cell Biol. 1976;71(3):848-75. PubMed PMID: 62755; PMCID: PMC2109793.

7. Bi E, Maddox P, Lew DJ, Salmon ED, McMillan JN, Yeh E, Pringle JR. Involvement of an actomyosin contractile ring in Saccharomyces cerevisiae cytokinesis. J Cell Biol. 1998;142(5):1301-12. PubMed PMID: 9732290; PMCID: PMC2149343.

8. Burki F, Roger AJ, Brown MW, Simpson AGB. The New Tree of Eukaryotes. Trends Ecol Evol. 2020;35(1):43–55. doi: 10.1016/j.tree.2019.08.008. PubMed PMID: 31606140.

9. Garcia-Salcedo JA, Perez-Morga D, Gijon P, Dilbeck V, Pays E, Nolan DP. A differential role for actin during the life cycle of Trypanosoma brucei. EMBO J. 2004;23(4):780–9. doi: 10.1038/sj.emboj.7600094. PubMed PMID: 14963487; PMCID: PMC381002.

10. Sherwin T, Gull K. The cell division cycle of Trypanosoma brucei brucei: timing of event markers and cytoskeletal modulations. Philos Trans R Soc Lond B Biol Sci. 1989;323(1218):573–88. doi: 10.1098/rstb.1989.0037. PubMed PMID: 2568647.

11. Wheeler RJ, Scheumann N, Wickstead B, Gull K, Vaughan S. Cytokinesis in Trypanosoma brucei differs between bloodstream and tsetse trypomastigote forms: implications for microtubule-based morphogenesis and mutant analysis. Mol Microbiol. 2013;90(6):1339–55. doi: 10.1111/mmi.12436. PubMed PMID: 24164479; PMCID: PMC4159584.

12. Robinson DR, Sherwin T, Ploubidou A, Byard EH, Gull K. Microtubule polarity and dynamics in the control of organelle positioning, segregation, and cytokinesis in the trypanosome cell cycle. J Cell Biol. 1995;128(6):1163-72. PubMed PMID: 7896879; PMCID: PMC2120423.

13. Hilton NA, Sladewski TE, Perry JA, Pataki Z, Sinclair-Davis AN, Muniz RS, Tran HL, Wurster JI, Seo J, de Graffenried CL. Identification of TOEFAZ1-interacting proteins reveals key regulators of Trypanosoma brucei cytokinesis. Mol Microbiol. 2018;109(3):306–26. doi: 10.1111/mmi.13986. PubMed PMID: 29781112; PMCID: PMC6359937.

14. Wickstead B, Gull K, Richards TA. Patterns of kinesin evolution reveal a complex ancestral eukaryote with a multifunctional cytoskeleton. BMC Evol Biol. 2010;10:110. doi: 10.1186/1471-2148-10-110. PubMed PMID: 20423470; PMCID: PMC2867816.

15. Wickstead B, Gull K. A “holistic” kinesin phylogeny reveals new kinesin families and predicts protein functions. Mol Biol Cell. 2006;17(4):1734–43. doi: 10.1091/mbc.E05-11-1090. PubMed PMID: 16481395; PMCID: PMC1415282.

16. Wozniak MJ, Milner R, Allan V. N-terminal kinesins: many and various. Traffic. 2004;5(6):400–10. doi: 10.1111/j.1600-0854.2004.00191.x. PubMed PMID: 15117314.

17. Vale RD, Milligan RA. The way things move: looking under the hood of molecular motor proteins. Science. 2000;288(5463):88–95. doi: 10.1126/science.288.5463.88. PubMed PMID: 10753125.

18. Hodges AR, Bookwalter CS, Krementsova EB, Trybus KM. A nonprocessive class V myosin drives cargo processively when a kinesin-related protein is a passenger. Curr Biol. 2009;19(24):2121–5. doi: 10.1016/j.cub.2009.10.069. PubMed PMID: 20005107; PMCID: PMC2904613.

19. Gheber L, Kuo SC, Hoyt MA. Motile properties of the kinesin-related Cin8p spindle motor extracted from Saccharomyces cerevisiae cells. J Biol Chem. 1999;274(14):9564–72. doi: 10.1074/jbc.274.14.9564. PubMed PMID: 10092642.

20. Lockhart A, Cross RA. Kinetics and motility of the Eg5 microtubule motor. Biochemistry. 1996;35(7):2365–73. doi: 10.1021/bi952318n. PubMed PMID: 8652578.

21. Matthies HJ, Baskin RJ, Hawley RS. Orphan kinesin NOD lacks motile properties but does possess a microtubule-stimulated ATPase activity. Mol Biol Cell. 2001;12(12):4000–12. doi: 10.1091/mbc.12.12.4000. PubMed PMID: 11739796; PMCID: PMC60771.

22. Muretta JM, Jun Y, Gross SP, Major J, Thomas DD, Rosenfeld SS. The structural kinetics of switch-1 and the neck linker explain the functions of kinesin-1 and Eg5. Proc Natl Acad Sci U S A. 2015;112(48):E6606–13. doi: 10.1073/pnas.1512305112. PubMed PMID: 26627252; PMCID: PMC4672802.

23. Parke CL, Wojcik EJ, Kim S, Worthylake DK. ATP hydrolysis in Eg5 kinesin involves a catalytic two-water mechanism. J Biol Chem. 2010;285(8):5859–67. doi: 10.1074/jbc.M109.071233. PubMed PMID: 20018897; PMCID: PMC2820811.

24. Kirchner J, Woehlke G, Schliwa M. Universal and unique features of kinesin motors: insights from a comparison of fungal and animal conventional kinesins. Biol Chem. 1999;380(7-8):915–21. doi: 10.1515/BC.1999.113. PubMed PMID: 10494842.

25. Edamatsu M. Molecular properties of the N-terminal extension of the fission yeast kinesin-5, Cut7. Genet Mol Res. 2016;15(1). doi: 10.4238/gmr.15017799. PubMed PMID: 26909973.

26. Zhernov I, Diez S, Braun M, Lansky Z. Intrinsically Disordered Domain of Kinesin-3 Kif14 Enables Unique Functional Diversity. Curr Biol. 2020. doi: 10.1016/j.cub.2020.06.039. PubMed PMID: 32649913.

27. Braun M, Drummond DR, Cross RA, McAinsh AD. The kinesin-14 Klp2 organizes microtubules into parallel bundles by an ATP-dependent sorting mechanism. Nat Cell Biol. 2009;11(6):724–30. doi: 10.1038/ncb1878. PubMed PMID: 19430466.

28. Sherwin T, Gull K. Visualization of detyrosination along single microtubules reveals novel mechanisms of assembly during cytoskeletal duplication in trypanosomes. Cell. 1989;57(2):211-21. PubMed PMID: 2649249.

29. Kashina AS, Baskin RJ, Cole DG, Wedaman KP, Saxton WM, Scholey JM. A bipolar kinesin. Nature. 1996;379(6562):270–2. doi: 10.1038/379270a0. PubMed PMID: 8538794; PMCID: PMC3203953.

30. Sharp DJ, McDonald KL, Brown HM, Matthies HJ, Walczak C, Vale RD, Mitchison TJ, Scholey JM. The bipolar kinesin, KLP61F, cross-links microtubules within interpolar microtubule bundles of Drosophila embryonic mitotic spindles. J Cell Biol. 1999;144(1):125–38. doi: 10.1083/jcb.144.1.125. PubMed PMID: 9885249; PMCID: PMC2148119.

31. Kapitein LC, Peterman EJ, Kwok BH, Kim JH, Kapoor TM, Schmidt CF. The bipolar mitotic kinesin Eg5 moves on both microtubules that it crosslinks. Nature. 2005;435(7038):114–8. doi: 10.1038/nature03503. PubMed PMID: 15875026.

32. McDonald HB, Stewart RJ, Goldstein LS. The kinesin-like ncd protein of Drosophila is a minus end-directed microtubule motor. Cell. 1990;63(6):1159–65. doi: 10.1016/0092-8674(90)90412-8. PubMed PMID: 2261638.

33. Carazo-Salas RE, Antony C, Nurse P. The kinesin Klp2 mediates polarization of interphase microtubules in fission yeast. Science. 2005;309(5732):297–300. doi: 10.1126/science.1113465. PubMed PMID: 16002618.

34. Carazo-Salas RE, Nurse P. Self-organization of interphase microtubule arrays in fission yeast. Nat Cell Biol. 2006;8(10):1102–7. doi: 10.1038/ncb1479. PubMed PMID: 16998477.

35. Daga RR, Lee KG, Bratman S, Salas-Pino S, Chang F. Self-organization of microtubule bundles in anucleate fission yeast cells. Nat Cell Biol. 2006;8(10):1108–13. doi: 10.1038/ncb1480. PubMed PMID: 16998476.

36. Janson ME, Loughlin R, Loiodice I, Fu C, Brunner D, Nedelec FJ, Tran PT. Crosslinkers and motors organize dynamic microtubules to form stable bipolar arrays in fission yeast. Cell. 2007;128(2):357–68. doi: 10.1016/j.cell.2006.12.030. PubMed PMID: 17254972.

37. Carleton M, Mao M, Biery M, Warrener P, Kim S, Buser C, Marshall CG, Fernandes C, Annis J, Linsley PS. RNA interference-mediated silencing of mitotic kinesin KIF14 disrupts cell cycle progression and induces cytokinesis failure. Mol Cell Biol. 2006;26(10):3853–63. doi: 10.1128/MCB.26.10.3853-3863.2006. PubMed PMID: 16648480; PMCID: PMC1488988.

38. Gruneberg U, Neef R, Li X, Chan EH, Chalamalasetty RB, Nigg EA, Barr FA. KIF14 and citron kinase act together to promote efficient cytokinesis. J Cell Biol. 2006;172(3):363–72. doi: 10.1083/jcb.200511061. PubMed PMID: 16431929; PMCID: PMC2063646.

39. Souto-Padron T, de Souza W, Heuser JE. Quick-freeze, deep-etch rotary replication of Trypanosoma cruzi and Herpetomonas megaseliae. J Cell Sci. 1984;69:167-78. PubMed PMID: 6386835.

40. Jones A, Faldas A, Foucher A, Hunt E, Tait A, Wastling JM, Turner CM. Visualisation and analysis of proteomic data from the procyclic form of Trypanosoma brucei. Proteomics. 2006;6(1):259–67. doi: 10.1002/pmic.200500119. PubMed PMID: 16302277.

41. Ohsugi M, Tokai-Nishizumi N, Shiroguchi K, Toyoshima YY, Inoue J, Yamamoto T. Cdc2-mediated phosphorylation of Kid controls its distribution to spindle and chromosomes. EMBO J. 2003;22(9):2091–103. doi: 10.1093/emboj/cdg208. PubMed PMID: 12727876; PMCID: PMC156080.

42. Yun M, Zhang X, Park CG, Park HW, Endow SA. A structural pathway for activation of the kinesin motor ATPase. EMBO J. 2001;20(11):2611–8. doi: 10.1093/emboj/20.11.2611. PubMed PMID: 11387196; PMCID: PMC125472.

43. Vale RD. Switches, latches, and amplifiers: common themes of G proteins and molecular motors. J Cell Biol. 1996;135(2):291-302. PubMed PMID: 8896589; PMCID: PMC2121043.

44. Bajar BT, Wang ES, Lam AJ, Kim BB, Jacobs CL, Howe ES, Davidson MW, Lin MZ, Chu J. Improving brightness and photostability of green and red fluorescent proteins for live cell imaging and FRET reporting. Sci Rep. 2016;6:20889. doi: 10.1038/srep20889. PubMed PMID: 26879144; PMCID: PMC4754705.

45. Howard J, Hyman AA. Preparation of marked microtubules for the assay of the polarity of microtubule-based motors by fluorescence microscopy. Methods Cell Biol. 1993;39:105-13. Epub 1993/01/01. PubMed PMID: 8246791.

46. Meijering E, Dzyubachyk O, Smal I. Methods for cell and particle tracking. Methods Enzymol. 2012;504:183–200. doi: 10.1016/B978-0-12-391857-4.00009-4. PubMed PMID: 22264535.

47. Schuck P. Size-distribution analysis of macromolecules by sedimentation velocity ultracentrifugation and lamm equation modeling. Biophys J. 2000;78(3):1606–19. doi: 10.1016/S0006-3495(00)76713-0. PubMed PMID: 10692345; PMCID: PMC1300758.

