## Supplemental Figures for "A *Trypanosoma brucei* orphan kinesin employs a convergent microtubule organization strategy to complete cytokinesis"

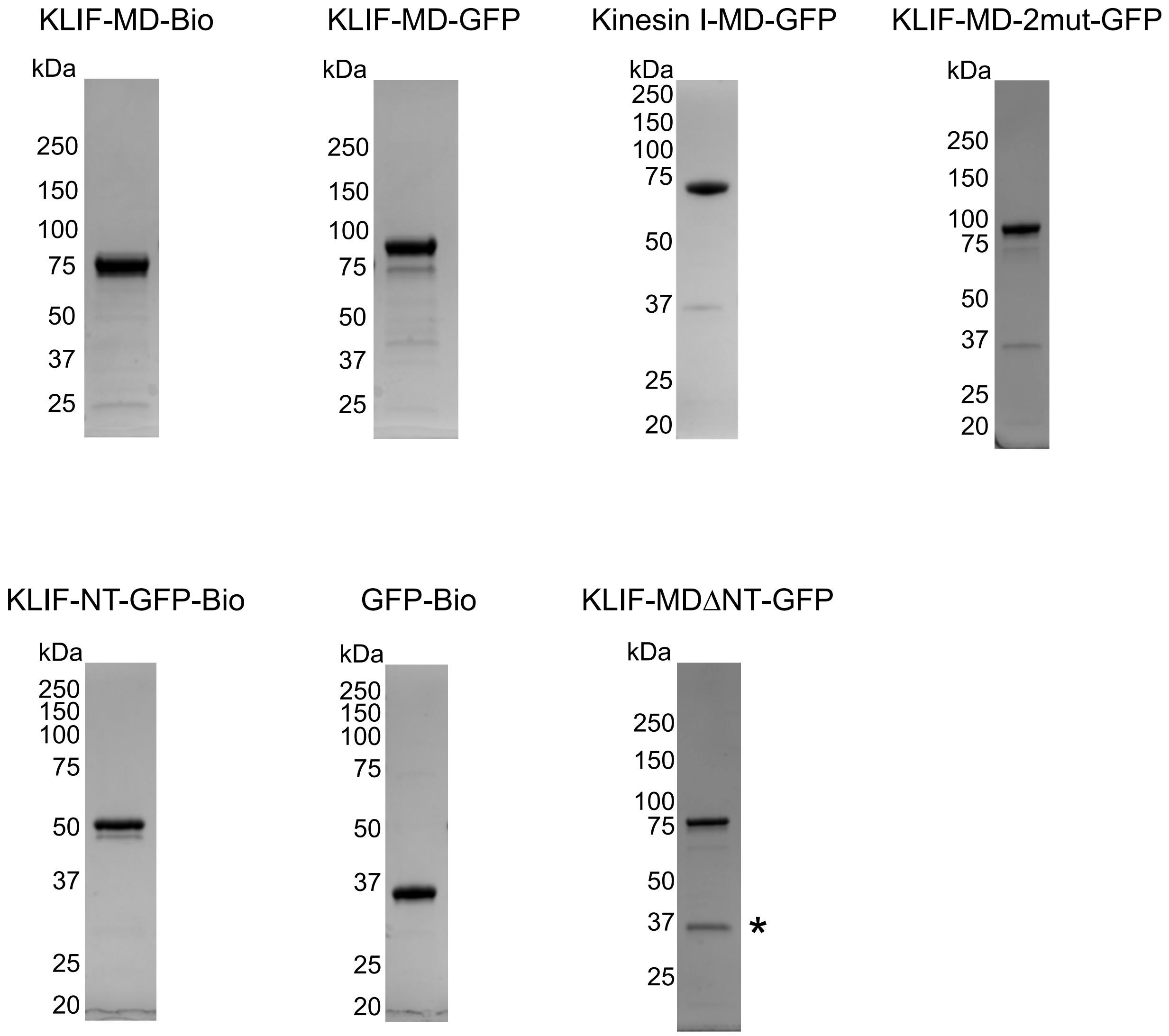


### Supplemental Figure S1

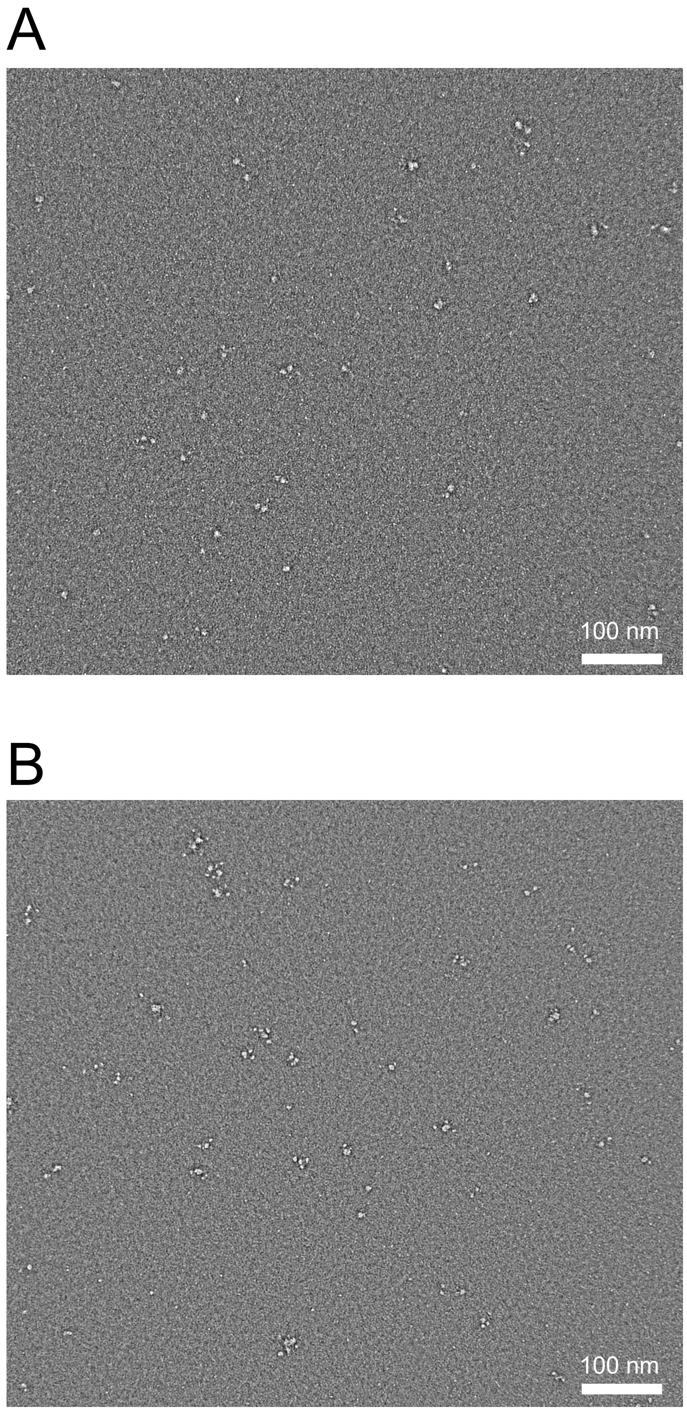


**Supplemental Figure S2**

A

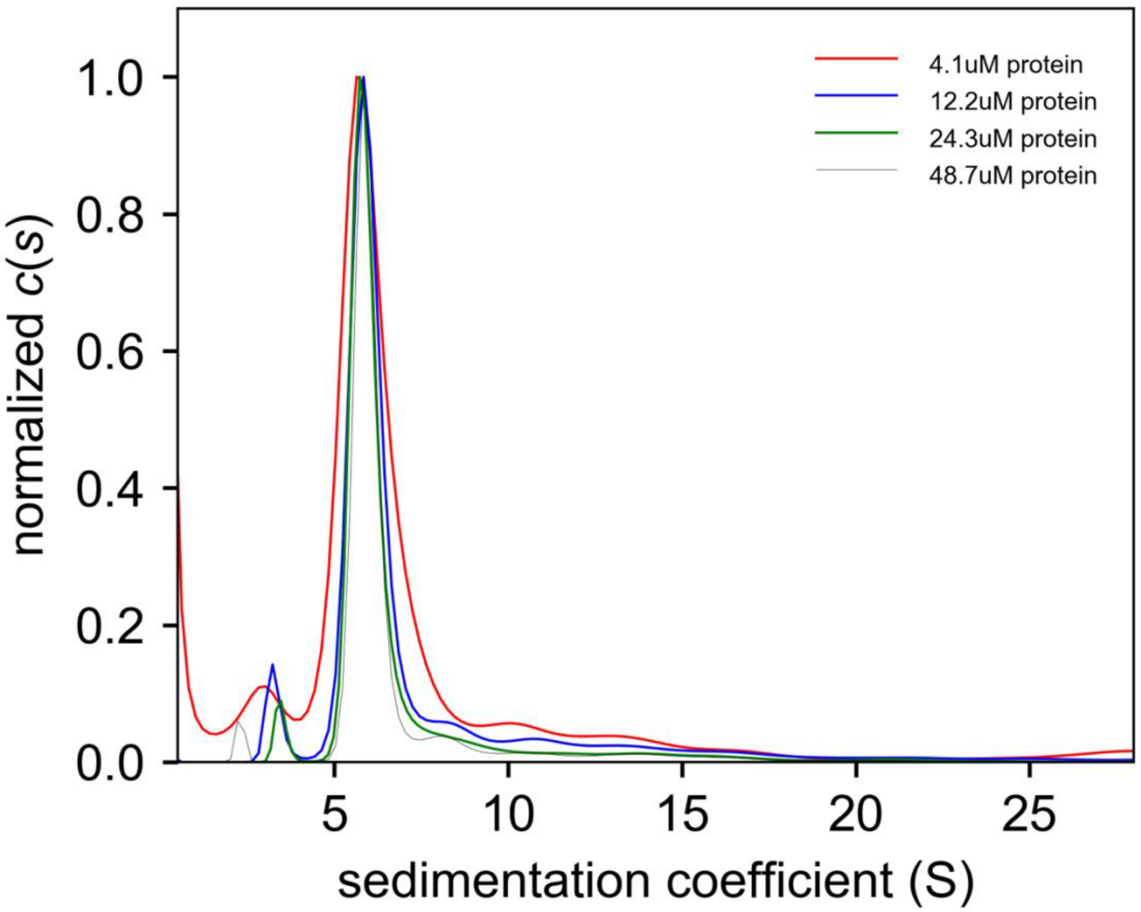


B

| Sample conc. (uM) | Peak(s) | | | | Frictional coefficient ($f/f_0$) | RMDS of fit |
|---|---|---|---|---|---|---|
| | 1 | 2 (Main dimer) | 3 | Aggr. | | |
| | S-value/MW (kDa)/% of tot.[&] | S-value/MW (kDa)/% of tot.[&] | S-value/MW (kDa)/% of tot.[&] | % of tot.[&] | | |
| 48.7 | 2.3S/36.6/2 | 6.0S/153.1/83 | 8.4S/256.6/6 | 9 | 1.64 (fitted) | 0.0032 |
| 24.3 | 3.5S/77.6/3 | 6.3S/189.7/91 | 15.1S/702.2/4 | 2 | 1.79 (fitted) | 0.0032 |
| 12.2 | 3.3S/69.3/5 | 6.2S/178.4/81 | 11.2S/432.6/6 | 8 | 1.74 (fitted) | 0.0026 |
| 4.1 | 2.9S/56.9/8 | 6.1S/172.7/78 | 14 | | 1.74 (fixed) | 0.0028 |

**&** Percent integrated area of all peaks, excluding peak at 0-0.3S which may be due to non-ideality or problems with fitting.

### Supplemental Figure S3

A

| | Switch 1 | | Switch 2 |
|---|---|---|---|
| KLIF | TAI**H**AR**SSRAH** | KLIF | LLA**DLAGCE**RIK |
| MmKIF2 | TSA**N**AH**SSRSH** | MmKIF2 | SLI**DLAGNE**RGA |
| DmNCD | TAG**N**ER**SSRSH** | DmNCD | NLV**DLAGSE**SPK |
| HsuKHC | TNM**N**EH**SSRSH** | HsuKHC | YLV**DLAGSE**KVS |
| MmKIF4 | TAM**N**SQ**SSRSH** | MmKIF4 | HLV**DLAGSE**RQK |
| MmKIF1A | TNM**N**ET**SSRSH** | MmKIF1A | SLV**DLAGSE**RAD |
| MmKIF13A | TNM**N**EE**SSRSH** | MmKIF13A | SLV**DLAGSE**RVS |
| Motif | NxxSSRSH | Motif | DxxGxE |
| | **A347** | | **A379** |

B

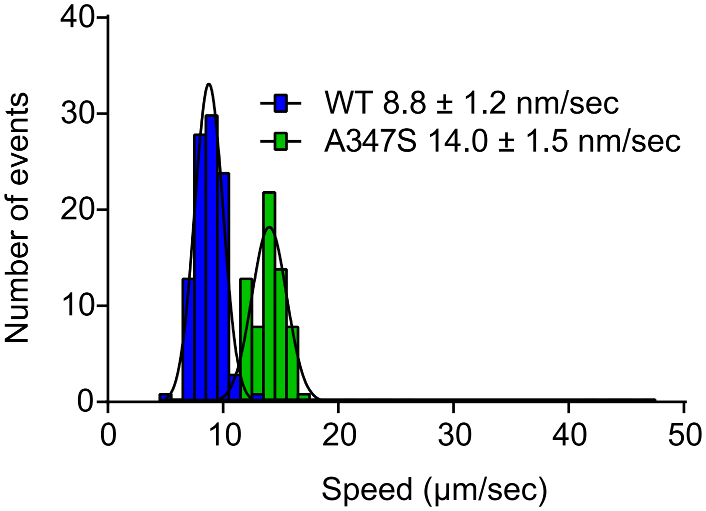


C

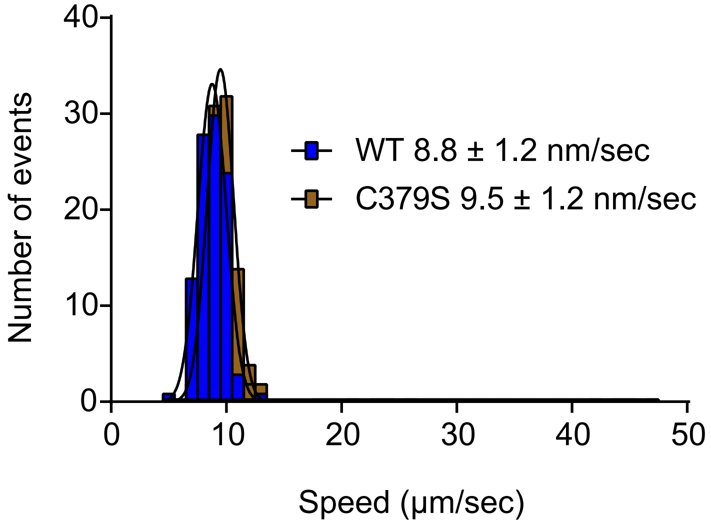


### Supplemental Figure S4

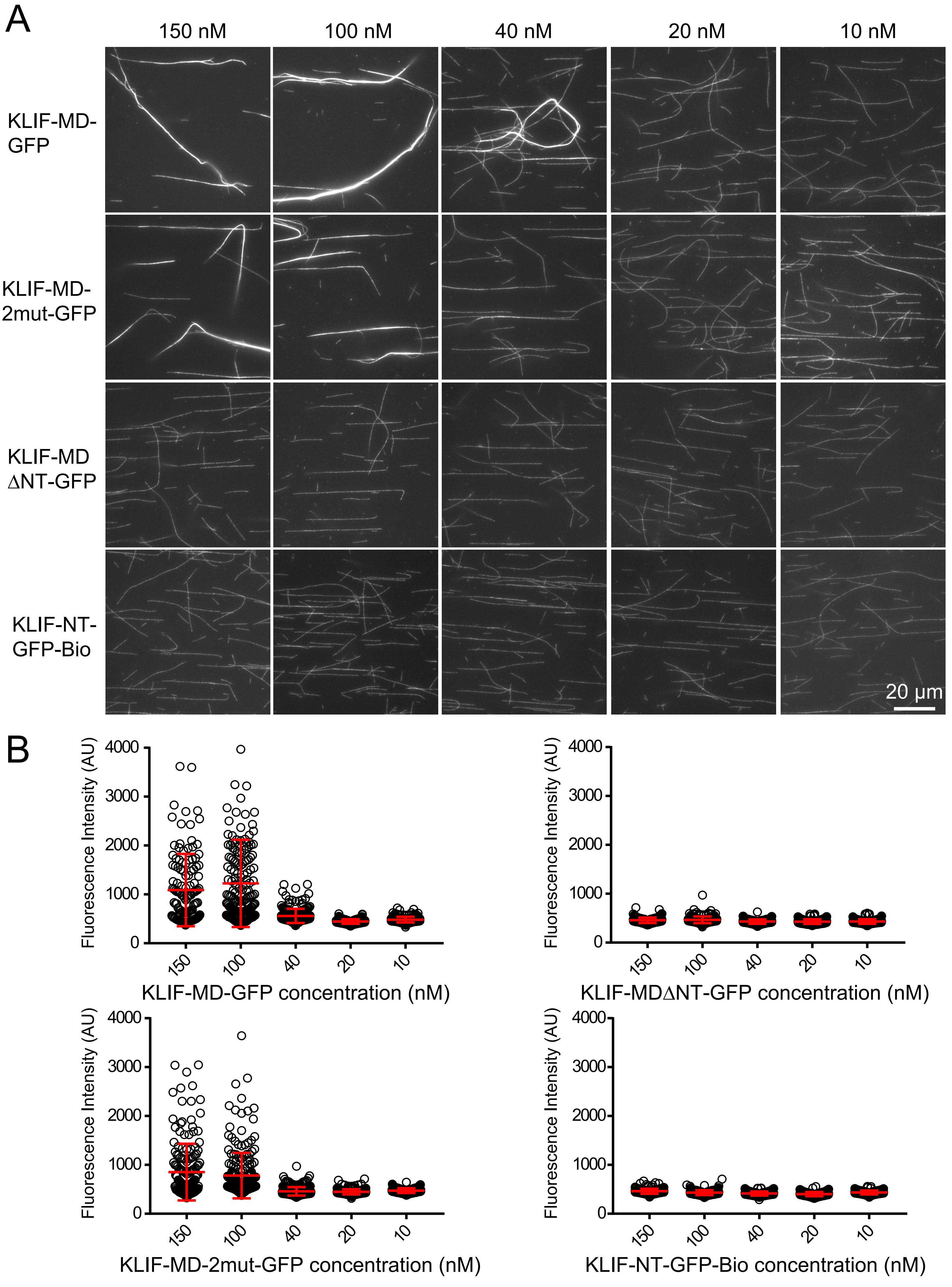


### Supplemental Figure S5
